# E2F1 Suppresses EBV Lytic Reactivation through Cellular and Viral Transcriptional Networks

**DOI:** 10.1101/2025.01.23.634435

**Authors:** Joyanta Biswas, SK Asif Ali, Samaresh Malik, Subhadeep Nag, Piyali Mukherjee, Abhik Saha

## Abstract

Latent EBV infection is causally associated with various B-cell malignancies, while periodic lytic-cycle replication is essential for sustaining viral progeny. Lytic cycle induction represents a promising therapeutic strategy for EBV-associated neoplasms. Therefore, uncovering the mechanisms that regulate EBV lytic-cycle reactivation is pivotal for understanding viral pathogenesis and advancing novel therapies. Our genome-wide transcriptomic analysis reveals that E2F1 expression is transcriptionally activated during EBV latent infection in B-lymphocytes but significantly suppressed during lytic cycle reactivation. While ectopic E2F1 expression suppresses lytic replication, E2F1 depletion markedly accelerates this process. Mechanistically, we establish that E2F1 and the lytic transactivator BZLF1 form a negative transcriptional feedback loop, tightly controlling viral lytic replication. Furthermore, E2F1 positively regulates c-Myc expression and together they repress the leaky BZLF1 expression during latency. Importantly, c-Myc does not influence E2F1 expression, nor does BZLF1 modulate c-Myc transcription, underlining a distinct regulatory hierarchy. In sum, our findings reveal that EBV tightly controls the latent-to-lytic switch through precise regulation of E2F1 expression, positioning E2F1 as a pivotal regulator of both cellular and viral gene expression.

**Synopsis:** EBV coordinates the latent-to-lytic switch by sensing E2F1 abundance, which acts as a crucial transcriptional regulator of both cellular and viral gene expressions.

- During EBV latent infection, E2F1 promotes c-Myc transcription, and together they suppress EBV lytic cycle transactivator BZLF1 expression.
- E2F1 and BZLF1 form a negative feedback loop in order to control each other’s transcriptions.
- BZLF1-driven controlled E2F1 expression successively inhibits c-Myc level, thereby stimulating EBV lytic cycle reactivation.
- BZLF1 does not regulate c-Myc, nor does c-Myc reciprocally regulate E2F1, emphasizing a unidirectional regulatory hierarchy.

## Introduction

Epstein-Barr virus (EBV), member of the gammaherpesvirus family, persistently infects more than 95% of the global population (Saha & Robertson, 2019; Young *et al*, 2016). Like any other herpesviruses, EBV similarly displays a biphasic life cycle comprising latent infection and lytic replication, both of which play crucial roles in its pathogenesis (Styles *et al*, 2017; Malik *et al*, 2024). While latent infection is associated with numerous cancers both of epithelial and lymphoid origins, EBV reactivation into lytic cycle replication is indispensable to maintain the production of progeny virus (Shannon-Lowe *et al*, 2017; Malik *et al*, 2024). A central feature of EBV latent infection is its ability to transform metabolically dormant B-lymphocytes into hyper-proliferating B-cell blasts, followed by the establishment of distinct latency programs (I, II, and III), each characterized by unique signatures of viral latent oncogene expression (Saha & Robertson, 2019; Burton & Gewurz, 2022). EBV employs these latency programs that ultimately direct B-cell blasts to enter germinal centre and differentiate into memory B-cells, which serve as the reservoir for lifelong infection (SoRelle *et al*, 2022). The restricted expression of EBV latent genes eases immune evasion and is a hallmark of its association with various B-cell malignancies, such as Burkitt’s lymphoma (BL), Hodgkin lymphoma (HL) and diffuse large B-cell lymphoma (DLBCL) (Shannon-Lowe *et al*, 2017). Studies reveal a significant connection between EBV-associated malignancies and the compromised host immune system, leading to post-transplant lymphoproliferative disorders (PTLDs) in immunosuppressive individuals, including organ transplant recipients and AIDS patients (Saha & Robertson, 2019; Shannon-Lowe *et al*, 2017).

In the germinal B-centre, infected memory B-cells can intermittently terminally differentiate into plasma cells and thereby triggering EBV lytic cycle reactivation, which is critical for sustaining the viral reservoir and promoting horizontal transmission from one host to another (Laichalk & Thorley-Lawson, 2005). This process involves re-expression of ∼80 viral genes required for active replication and the production of viral progeny (Yap *et al*, 2022). EBV lytic gene expression occurs in a temporally regulated cascade, which can be broadly divided into four phases - immediate-early (IE), early (E), leaky late (LL), and late (L). The lytic cycle replication is began by transcriptional activation of one or both of its IE promoters - Zp and Rp, leading to expression of two IE genes, BZLF1 (encoding Zta) and BRLF1 (encoding Rta), respectively (Yap *et al*, 2022; Wang *et al*, 2024). Although both are transcription factors (TFs) and can transactivate each other’s promoters to initiate viral reactivation, in most of the EBV positive cells BZLF1 can solely orchestrate EBV lytic cycle replication by provoking transcriptional activation of nearly 30 E genes (Bernaudat *et al*, 2022; Guo *et al*, 2020a). Newly replicated viral DNA then serve as templates for further transcriptions of L lytic genes. Some L-phase genes, referred to as LL, may begin expressing before completing entire viral genome replication. Expression of these genes facilitate encapsidation of viral dsDNA, virion assembly at the cell membrane, and egression of mature infectious particles (Yap *et al*, 2022; Wang *et al*, 2024; Guo *et al*, 2020a). In contrast to latent genes, EBV lytic antigens are highly immunogenic, eliciting strong immune responses. However, the temporal regulation of lytic genes ensures efficient viral replication by minimizing immune detection (Taylor *et al*, 2015).

As yet, there are no clinically approved therapies available that selectively targets EBV associated cancers. Induction of EBV lytic cycle followed by treatment with antiviral drugs offers an attractive therapeutic approach to treat EBV associated cancers (Yiu *et al*, 2020). This targeted ‘lytic induction therapy’ functions by triggering lytic cycle replication, during which two viral protein kinases BXLF1 and BGLF4 phosphorylate nucleoside analogues like acyclovir and ganciclovir, converting the pro-drugs into their active nucleotide forms, resulting in cytotoxicity in both EBV-infected and neighbouring cells (Meng *et al*, 2010). Therefore, there is increasing interest in strategies that drive EBV reactivation from latency into the lytic cycle, thereby sensitizing EBV-positive tumour cells to these nucleoside analogues. However, the existing methods of EBV lytic cycle reactivation by chemical inducers are highly cytotoxic and seldom lytic replication occurs in only a small percentage of latently infected cells (Malik *et al*, 2024). To date, among the growing list of EBV lytic cycle inducers, only a few have been tested in clinical trials, but failed due to high cytotoxicity and incongruous pharmacokinetics (Yiu *et al*, 2020). A proper understanding of cell mechanisms controlling the balance between latent infection and lytic replication is therefore critical for further development of potential drugs that selectively kill EBV positive tumour cells. EBV latent-to-lytic switch is tightly controlled by several cellular factors, including TFs, chromatin remodelling, and signalling pathways (Wang *et al*, 2024). For example, during B-cell differentiation into plasma cells, BLIMP1/PRDM1 and XBP-1 TFs induce spontaneous EBV lytic replication by transcriptional activation of both IE genes - BZLF1 and BRLF1 (Bhende *et al*, 2007; Reusch *et al*, 2014). Recently, genome-wide CRISPR-Cas9 screening in EBV-positive BL cells identified c-Myc TF as a negative regulator of lytic cycle replication (Guo *et al*, 2020a). Our previous global transcriptomic analysis revealed that E2F1 expression is transcriptionally activated during EBV latent infection in naïve B-lymphocytes but suppressed during reactivation into lytic-cycle replication (Malik *et al*, 2024; Gain *et al*, 2020). E2F1, the first member of E2F family TFs, plays a central role in cell cycle progression, particularly the transition from the G1 to S phase, as well as DNA-damage response and apoptosis (Choi & Kim, 2019). Accumulating evidence indicates that E2F1 regulates replication of several DNA viruses (Xiaofei *et al*, 2011; Johnson *et al*, 2002). Although E2F1 functions are implicated in EBV pathogenesis, particularly in latently infected cells (Saha *et al*, 2012; Boreström *et al*, 2012; Pei *et al*, 2016), there are no reports describing its role in lytic cycle replication. Given that E2F activity is often deregulated by infection with DNA viruses including EBV, we hypothesize that E2F1 contributes to EBV induced B-cell lymphomagenesis by regulating its latent-to-lytic switch. Overall our data support a model in which E2F1 transcriptionally controls both cellular (c-Myc) and viral (BZLF1) gene expressions to suppress EBV lytic replication.

## Results

### EBV reactivation to lytic cycle replication suppresses E2F1 expression

The E2F family of TFs is crucial for regulating cell cycle progression, DNA replication and apoptosis. In mammalian cells, there are eight E2F genes encoding 10 different isoforms, each has distinct functions in cell fate. The E2F TFs can be categorized as transcription activators (E2F1-3), canonical repressors (E2F4-6) and non-canonical repressors (E2F7-8). Among the E2F isoforms, elevated E2F1 expression is typically implicated with the poor prognosis of several solid cancers. Accumulating evidence also suggests a direct correlation of E2F1 activities on EBV induced B-cell lymphomagenesis. Reanalysis of our previous RNA sequencing (RNA-Seq) data (GSE235941) of EBV latently infected naïve B-lymphocytes (0-4 days post-infection, dpi) as well as microarray data of EBV infected BL line BL31 revealed significant transcriptional activation of E2F1 along with three other E2F isoforms E2F2, E2F7 and E2F8 (Fig EV1A-B). A similar trend of transcriptional activation of E2F1 was also witnessed in reanalysis of RNA-Seq data (GSE125974) and qRT-PCR of EBV induced B-cell transformation (0-28 dpi) (Fig EV1C-D). RNA-Seq data of ‘Genotype-Tissue Expression (GTEx; https://gtexportal.org/home/) portal and qRT-PCR analysis further substantiated E2F1 transcriptional activation in EBV transformed lymphoblastoid cell lines (LCLs) (Fig EV1E-F).

In contrast to latent infection, there are no or contradictory reports available on E2F1 regulations during EBV lytic replication. Elevated expressions of specific set of E2F isoforms during EBV latent infection prompted us to further investigate expression pattern of these E2F members during lytic cycle reactivation. Various small molecules have been identified as stimulators of EBV lytic cycle replication (Yiu *et al*, 2020). Combination of TPA (12-O-tetradecanoylphorbol-13-acetate) and sodium butyrate (NaBu) are being largely utilized in laboratory settings to induce EBV reactivation into lytic cycle replication from EBV positive B-cells. Recently our lab has also demonstrated that proteasomal inhibition promotes EBV lytic cycle replication (Gain *et al*, 2020). Reanalysis of our previous RNA-Seq data of both TPA /NaBu (GSE237484; Malik *et al*, 2024) and proteasomal inhibition by MG132 (Gain *et al*, 2020) induced EBV lytic cycle reactivation in LCLs demonstrated significant transcriptional repression of these specific E2F isoforms – E2F1, E2F2, E2F7 and E2F8 (Fig EV1G-H).

Based on its impact on cell cycle and apoptosis as well as its well-known oncogenic properties (Kumari *et al*, 2015; Shats *et al*, 2017), we further focused our study unequivocally on E2F1. In order to rule out inconsistencies due to chemically induced reactivation of EBV lytic cycle replication, several methods of lytic cycle induction was opted including treatment with combination of TPA/NaBu, B-cell receptor (BCR) activation by IgG crosslinking or proteasomal inhibition by MG132 in multiple EBV positive cells (Fig 1). Expressions of EBV encoded IE gene BZLF1 and E gene EaD/BMRF1 by immunoblot and qRT-PCR analysis were employed to determine lytic cycle reactivation (Fig 1). As similar to RNA-Seq data, TPA/NaBu treatment mediated EBV reactivation led to significant downregulation of E2F1 expressions in both LCLs and patient-derived BL line EB3 (Fig 1A-D). To understand whether EBV lytic replication in epithelial background also equally represses E2F1 expression, we utilized HEK293T cells stably harbouring GFP-tagged EBV bacmid (Fig 1E-G). Parallel to EBV positive B-lymphocytes, TPA/NaBu treatment equally transcriptionally suppressed E2F1 expression during viral lytic cycle reactivation (Fig 1E-G). Of the several methods used in the laboratory settings for EBV lytic cycle induction, BCR activation represents the most physiologic. Antigen recognition by the BCR can be recapitulated *in vitro* by cross-linking of surface immunoglobulins - IgG or IgM, depending upon which immunoglobulin is produced by the B-cell line. IgG treatment for 72 h of another patient-derived EBV positive BL line P3HR1 revealed similar E2F1 depletion (Fig 1H). In addition, proteasomal inhibition by MG132 treatment also exhibited similar transcriptional repression in both HEK293T cells harbouring GFP-EBV bacmid and LCL#89 (Fig 1I-K). Overall, these results evidently demonstrated significant transcriptional repression of E2F1 in response to EBV lytic cycle reactivation in both epithelial and B-cell background, raising the question of whether EBV adjusts E2F1 expression in order to sensitize latently infected cells for lytic cycle induction.

**Figure 1.**
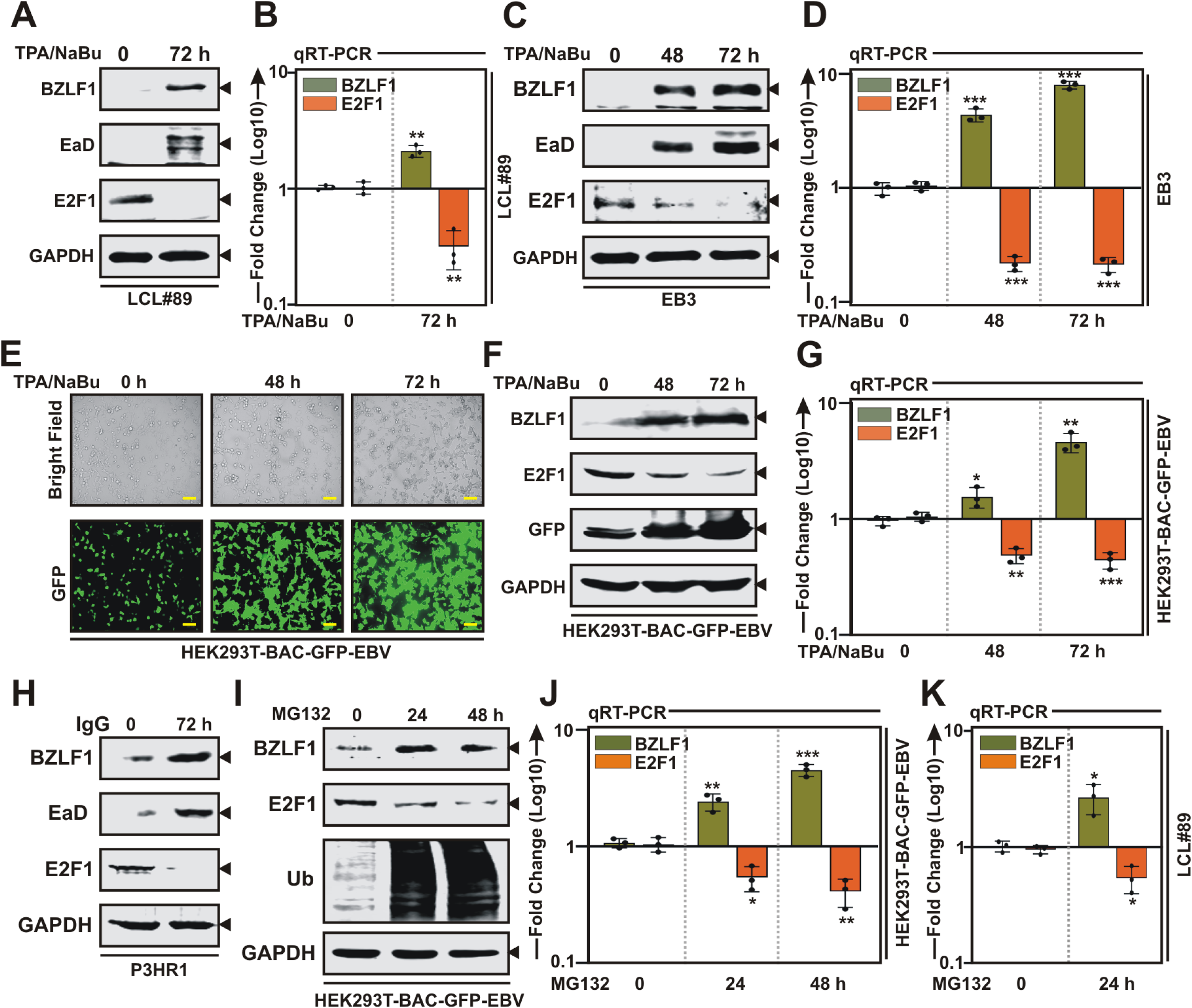
EBV lytic reactivation in both B- and epithelial cells depletes E2F1 expression. **(A-K)** EBV lytic cycle replication was induced by with 20 ng/ml 12-O-tetradecanoylphorbol-13-acetate (TPA) and 3 mM sodium butyrate (NaBu) treatment in **(A-B)** EBV transformed B-lymphocytes LCL#89, **(C-D)** EBV^+^ BL line EB3, **(E-G)** HEK293T cells harbouring GFP-EBV Bacmid for the indicated time points (0-72 h) and subjected to **(A, C, F)** immunoblot and **(B, D, G)** qRT-PCR analysis. **(E)** Representative pictures of bright field and GFP fluorescence of TPA/NaBu treated HEK293T-BAC-GFP-EBV cells. Scale bars, 100 μm. **(H)** EBV lytic cycle replication was induced in EBV^+^ BL line P3HR1 with anti-human IgG (10 µg/ml) for 72 h and subjected for immunoblot analysis. **(I-J)** HEK293T-BAC-GFP-EBV or **(K)** LCL#89 either left untreated or treated with 1 μM MG132 for the indicated hours and subjected to **(I)** immunoblot or **(J-K)** qRT-PCR analysis. **(A, C, F, H, I)** Blots are representative of n = 3 replicates. **(B, D, G, J, K)** For qRT-PCR analysis, the relative changes in transcripts of the selected genes were quantified using the 2^−ΔΔCT^ method and normalized with B2M as internal control. The results are presented as the mean ± SD, n = 3 biological replicates. Statistical significance was determined by a two-sided Student’s t-test, *P < 0.05; **P < 0.01; ***P < 0.001; ns, not significant. Source data are available online for this figure.

### EBV immediate early protein BZLF1 transcriptionally represses E2F1 expression

To understand the underlying molecular mechanism governing E2F1 transcriptional activation during EBV driven B-cell transformation or virally transformed B-lymphocytes, we first reanalyzed chromatin immunoprecipitation followed by deep sequencing (ChIP-Seq) data for all six EBV nuclear antigens (EBNAs) - EBNA1 (GSE73887), EBNA2 (GSE29498), EBNALP (GSE49338), and three EBNA3 genes (GSE88729) - EBNA3A, EBNA3B and EBNA3C in LCLs (Fig S1 and Table EV1). Additionally, since EBV encoded major latent membrane protein, LMP1 constitutively activates NF-κB signalling to promote LCLs growth and survival (Lavorgna & Harhaj, 2012), we included ChIP-Seq data (GSE55105) for all NF-κB subunits RelA, RelB, cRel, p50, and p52 in our analysis (Fig S1 and Table EV1). Reanalysis of ChIP-Seq data revealed no viral genes along with NF-κB subunits were enriched in the E2F1 promoter region (Fig S1). Non-occupancy of latent viral oncoproteins coupled with transcriptional repression during lytic cycle reactivation prompted us to further investigate the binding capacity of EBV lytic proteins on E2F1 promoter region. BZLF1 belongs to the family of basic leucine zipper (bZIP) TFs, may solely induce EBV lytic cycle replication (Bernaudat *et al*, 2022). In addition to regulating viral lytic gene transcriptions, BZLF1 can also transcriptionally regulate expressions of multiple cell-cycle genes (Mauser *et al*, 2002). Moreover, there are two contradictory reports demonstrating while BZLF1 can induce E2F1 expression in primary keratinocytes and gastric carcinoma cells (Mauser *et al*, 2002), E2F1 along with c-Myc obstruct BZLF1 mediated transactivation through a negative regulatory element located at the N-terminal region (Lin *et al*, 2004).

Reanalysis of ChIP-Seq data (E-MTAB-7788) of Raji cells stably expressing BZLF1 under doxycycline responsive promoter revealed several distinct BZLF1 ChIP-Seq signals in the E2F1 promoter region (Fig 2A). Model-based analysis of ChIP-Seq (MACS) tool for peak-calling with a p-value cut off set at p<0.05 up to ∼4 Kb upstream of the TSS (transcription start site) identified significant BZLF1 peak on the E2F1 promoter region (Fig 2A). BZLF1 is related to cellular AP-1 (activating protein 1) family of TFs that binds to two different classes of BZLF1-responsive elements on DNA - the canonical AP-1 binding site (TGACTCA) and methylated CpG containing motifs. The identified ChIP-Seq signal on E2F1 promoter were assessed for BZLF1/AP-1 binding motifs using JASPAR tool and subsequently revealed three AP-1 motifs (Fig 2A-B). BZLF1 occupancy on E2F1 promoter was validated by ChIP-qPCR using P3HR1 cells with or without EBV lytic cycle induction (Fig 2C). As compared to the control there was a significant enrichment of BZLF1 binding onto E2F1 promoter region upon lytic cycle reactivation (Fig 2C). Lytic cycle reactivation in P3HR1 cells using a similar experimental set up also resulted in significant downregulation of E2F1 expression at protein level (Fig 2D).

**Figure 2.**
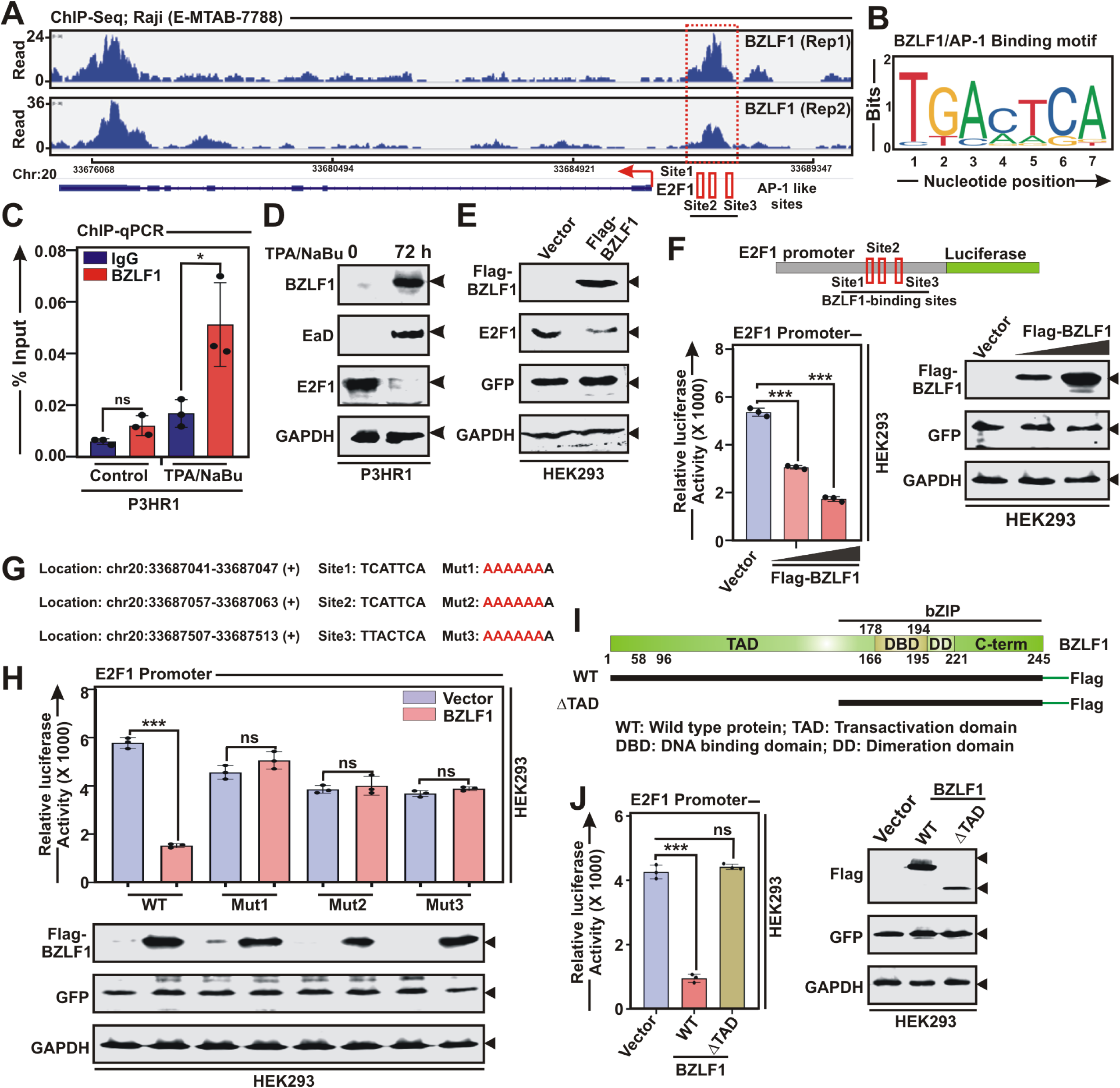
BZLF1 transcriptionally represses E2F1 expression. **(A)** Reanalysis of ChIP-Seq data (E-MTAB-7788) showing enrichment of BZLF1 on E2F1 promoter. Bottom panel indicates the MACS identified peaks (Sites 1-3) for BZLF1/AP-1 binding on E2F1 promoter. **(B)** BZLF1 homologue AP-1 binding motif identified on the MACS peaks of E2F1 promoter region. **(C)** ChIP-qPCR data showing recruitment of BZLF1 on E2F1 promoter upon EBV lytic cycle reactivation by TPA/sodium butyrate (NaBu) for 72 h in P3HR1. **(D)** P3HR1 reactivated to lytic cycle replication as similar to **(C)** were subjected to immunoblot analysis. **(E)** HEK293 cells transiently transfected with empty vector or flag-tagged BZLF1 expression plasmid for 36 h were harvested and subjected to immune blot analysis. **(F)** Luciferase reporter activity and the corresponding immunoblot analysis of the wild-type E2F1 promoter in the presence of increasing concentrations of BZLF1 expression plasmid in transiently transfected HEK293 cells. **(G)** Schema showing three wild-type BZLF1/AP-1 binding sites (Sites 1-3) and their corresponding mutations (Muts 1-3) on E2F1 promoter for cloning into pGL3 luciferase reporter vector. **(H)** Luciferase reporter activity of the wild-type and the mutant E2F1 promoters in the presence of either vector control or BZLF1 expression plasmid. A fraction of the total protein were evaluated by immunoblot analysis. **(I)** Schema showing different structural domains of BZLF1 for cloning in flag-tagged expression vector. **(J)** Luciferase reporter activity and the corresponding immunoblot analysis of the E2F1 promoter in the presence of empty vector, wild-type (WT) or transactivation domain deleted (ΔTAD) BZLF1 expression plasmids. The results are presented as the mean ± SD, n = 3 biological replicates. Statistical significance was determined by a two-sided Student’s t-test, *P < 0.05; **P < 0.01; ***P < 0.001; ns, not significant. Source data are available online for this figure.

Next, we wanted to check whether BZLF1 induction during lytic cycle replication is directly accountable for E2F1 transcriptional repression. HEK293 cells transiently transfected with flag-tagged BZLF1 expression vector demonstrated that BZLF1 expression was inversely correlated with the endogenous E2F1 expression (Fig 2E). To rule out BZLF1 mediated post-translational activities, HEK293 cells were further transiently transfected with myc-tagged BZLF1 with or without flag-tagged E2F1 expression vectors and subjected to immunoblot and co-immunoprecipitaion analysis (Fig S2A-B). In contrast to endogenous E2F1, BZLF1 had effect no effect on exogenously expressed flag-E2F1 (Fig S2A), nor these two proteins interacted with each other (Fig S2B), suggesting the possibility of BZLF1 mediated transcriptional repression of E2F1 in EBV positive cells during lytic cycle replication. BZLF1 mediated depletion of E2F1 expression in transiently transfected HEK293 cells was further assessed by luciferase based promoter assay (Fig 2F). E2F1 promoter region comprising of three BZLF1/AP-1 binding sites were inserted upstream of the luciferase gene in pGL3-basic vector (Fig 2F). BZLF1 significantly repressed E2F1 promoter activity in a dose dependent manner (Fig 2F). Upon mutation of all three BZLF1/AP-1 sites on the E2F1 promoter region, BZLF1 failed to repress E2F1 transcription (Fig 2G-H), further validating BZLF1 mediated transcriptional suppression of E2F1 activity. We and others previously demonstrated that while BZLF1 N-terminal transactivation domain (TAD) controls gene transcription, the C-terminal basic leucine-zipper (bZIP) domain exhibits comparable DNA binding activity as the wild-type (WT) protein. In agreement with this our results also demonstrated unlike WT BZLF1, ΔTAD alone failed to repress E2F1 promoter activity (Fig 2I-J). Collectively, these results are consistent with a model in which upon lytic cycle induction EBV encodes its first lytic protein BZLF1 that in turn transcriptionally represses E2F1 expression.

### E2F1 negatively regulates BZLF1 expression

Given its role as a robust DNA-binding transcription factor and the observed differential expressions between EBV latent infection and lytic cycle reactivation, we hypothesized that E2F1 directly interacts with the EBV genome to regulate viral gene transcription. We employed ChIP-Seq to assess E2F1 binding across the EBV genome in two LCLs (Fig 3A). E2F1 occupancy was observed near all the EBV latent promoters – Wp, Cp, Qp, LMP1p and LMP2p, as confirmed by ChIP-qPCR analysis in both LCLs (Fig 3B). Within the lytic replication genomic regions significant E2F1 occupancy was only observed at the BZLF1 promoter/Zp (Fig 3B). However, no E2F1 enrichment was observed near both OriLyt regions – OriLytL and OriLytR (Fig 3B). BZLF1 directly binds to the OriLyt region and initiate EBV lytic genome replication. Recently, c-Myc was shown to directly bind to OriLyt region, thereby controlling BZLF1 occupancy as well as EBV latent-to-lytic switch (Guo et al, 2020a). To further pinpoint the putative binding sites of E2F1 within the proximal genomic region to the ChIP-Seq signals, JASPAR web tool (Rauluseviciute *et al*, 2024) was utilized. We identified multiple putative E2F1 binding motifs (GGCGGGAAA) near all the viral promoters including Zp (Fig 3C and Table EV2). We independently cloned these EBV genomic regions in pGL3-basic vector and performed promoter assays in the absence and presence of E2F1 (Figs 3D-E and EV2). Among the latent promoters, varied E2F1 regulations were observed (Fig EV2). While E2F1 transcriptionally repressed Wp, Cp and both LMP1p and LMP2p activities, positive transcriptional regulation was observed for Qp (Fig EV2). Notably, E2F1 transcriptionally repressed Zp activity in a dose dependent fashion (Fig 3E). Zp contains three distinct E2F1 binding sites (sites 1–3), and combinatorial mutational analysis revealed that while all three are important, site 2 is indispensable for E2F1-mediated transcriptional repression (Fig 3E).

**Figure 3.**
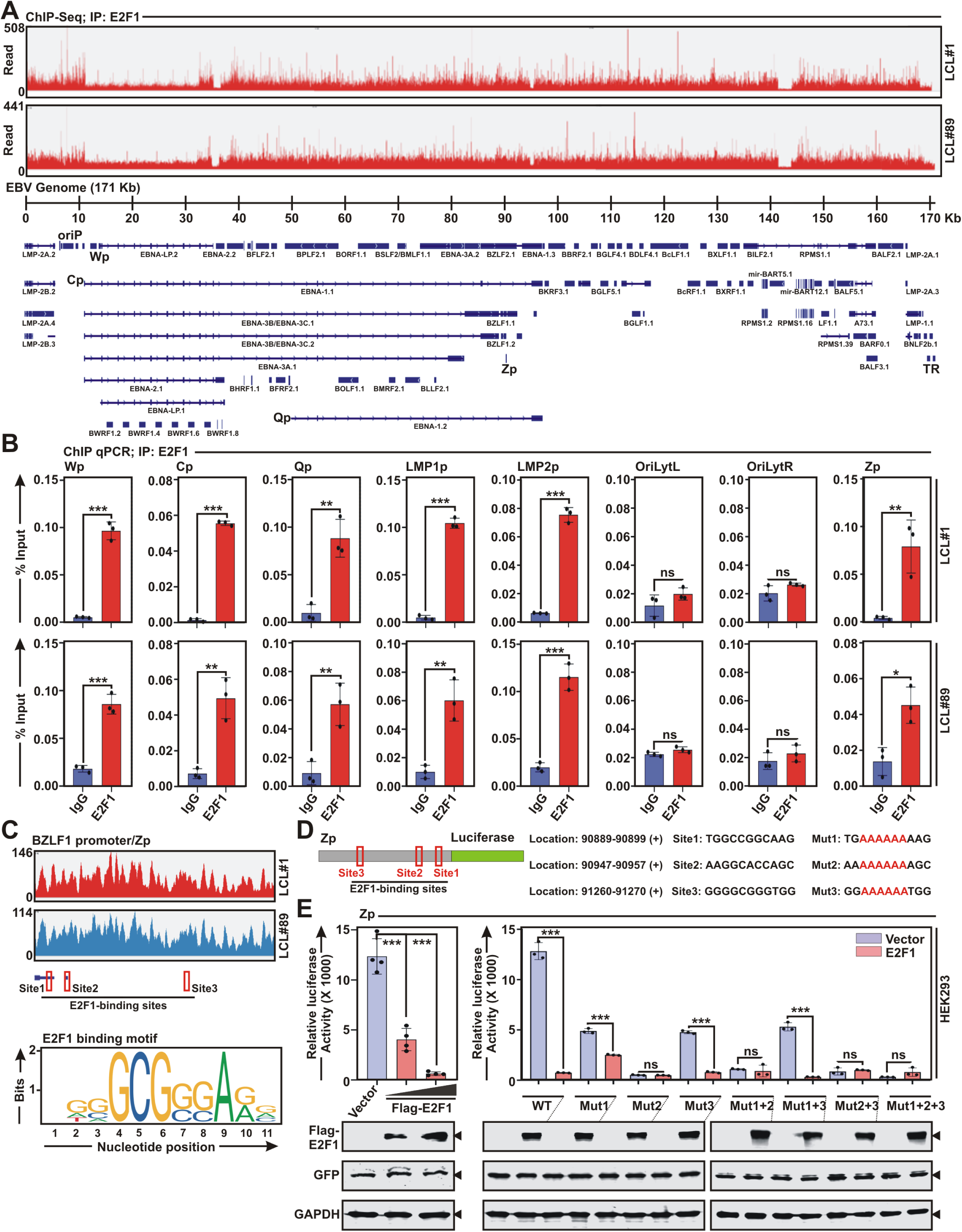
E2F1 negatively regulates BZLF1 expression. **(A)** ChIP-Seq analysis of E2F1 binding in two EBV transformed lymphoblastoid cell lines – LCL#1 and LCL#89. Tracks are aligned with the annotated EBV genome shown at the bottom. **(B)** ChIP-qPCR analysis of E2F1 occupancy at different EBV promoters and genomic regions. Anti-E2F1 ChIP was performed on chromatin extracted from LCL#1 and LCL#89, followed by qPCR using primers specific for Wp, Cp, Qp, LMP1p, LMP2p, OriLytL, OriLytR and Zp regions. Data are presented as % input. **(C)** E2F1 ChIP-Seq tracks on BZLF1 promoter region (Zp) in LCL#1 and LCL#89. Bottom panel indicates three putative E2F1 binding motifs obtained from JASPAR database in Zp. **(D)** Schema showing three wild-type E2F1 binding sites (Sites 1-3) and their corresponding mutations (Muts 1-3) on Zp for cloning into pGL3 luciferase reporter vector. **(E)** Luciferase reporter activity and the corresponding immunoblot analysis of the wild-type (WT) and mutant (Mut) Zp in the absence and presence of flag-tagged E2F1 expression vector. The results are presented as the mean ± SD, n = 3 biological replicates. Statistical significance was determined by a two-sided Student’s t-test, *P < 0.05; **P < 0.01; ***P < 0.001; ns, not significant. Source data are available online for this figure.

### E2F1 expression controls EBV lytic cycle reactivation

Hitherto, our data demonstrated that EBV lytic cycle transactivator BZLF1 and E2F1 negatively cross-regulate each other to maintain either latent or lytic replication status. We hypothesized that E2F1 expression might control EBV latent-to-lytic switch and lessening E2F1 levels could promote viral lytic replication. We first checked E2F1 expressions in several EBV positive BL lines – P3HR1, Jiyoye, EB3 and Namalwa (Fig 4A). While P3HR1, EB3 and Namalwa had significant E2F1 expressions, Jiyoye exhibited little or negligible E2F1 expression (Fig 4A). To investigate the role of E2F1 in regulating EBV lytic cycle replication, we further utilized P3HR1 and Jiyoye cells with distinct expression pattern of E2F1. Notably, both P3HR1 and Jiyoye contain type 2 EBV DNA, making them suitable for comparison in experimental setups. Upon lytic cycle reactivation, Jiyoye with little or no E2F1 expression significantly enhanced viral lytic cycle replication as compared to P3HR1 with elevated E2F1 expression (Fig 4B-D). The results were evaluated by immunoblot and qRT-PCR analysis of viral lytic gene expressions – BZLF1 and BMRF1/EaD (Fig 4B-C) as well as by quantifying both intracellular and extracellular EBV genome copy number (Fig 4D).

**Figure 4.**
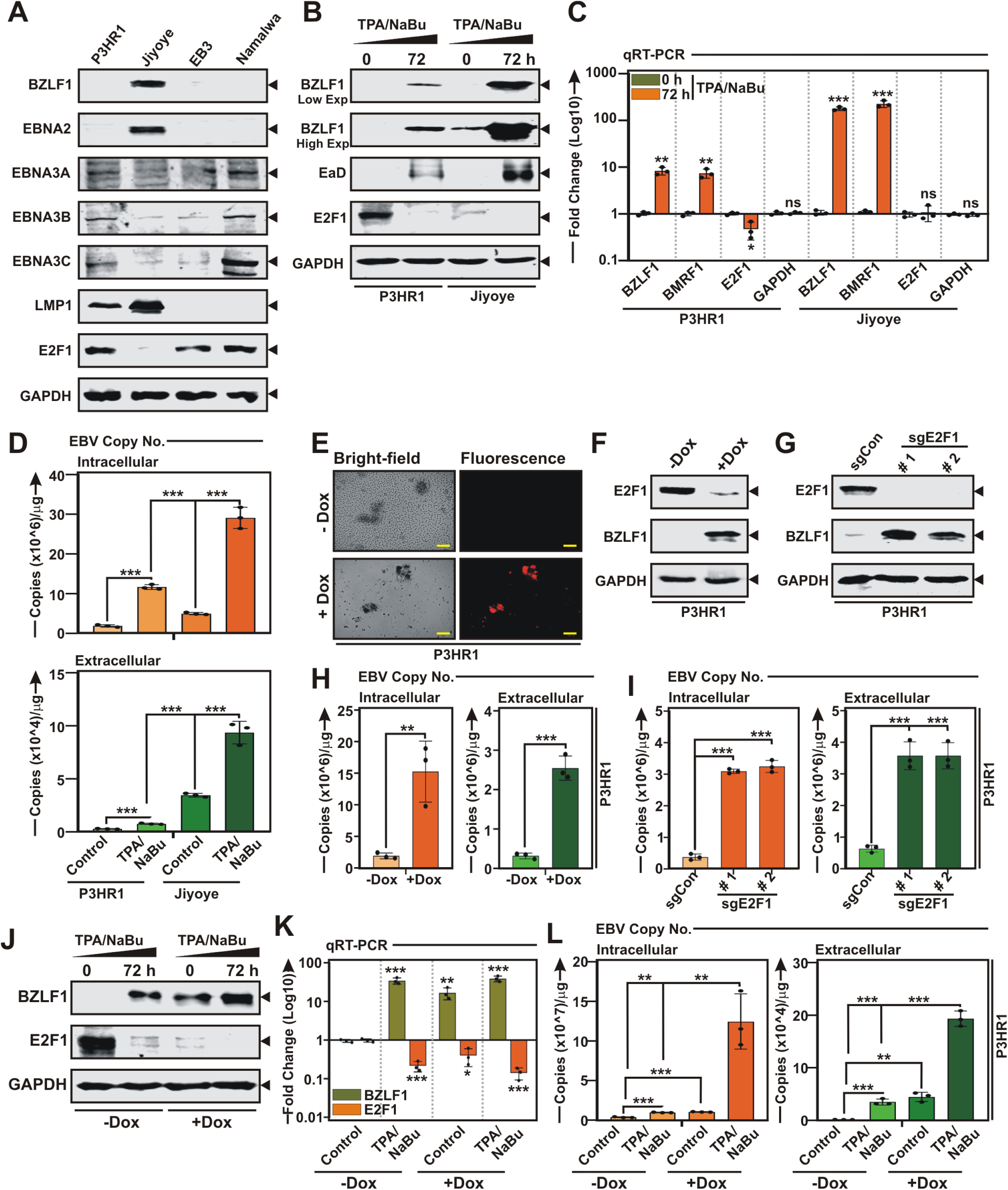
E2F1 depletion triggers EBV lytic cycle reactivation. **(A)** Immunoblot analysis of whole cell extracts of EBV^+^ BL lines – P3HR1, Jiyoye, EB3 and Namalwa with the indicated antibodies against viral and cellular proteins. **(B-C)** P3HR1 and Jiyoye cells reactivated to lytic cycle replication by TPA/sodium butyrate (NaBu) treatment for 72 h were either subjected to **(B)** immunoblot or **(C)** qRT-PCR analysis. **(D)** qRT-PCR of EBV intracellular or DNase-treated extracellular genome copy number from P3HR1 and Jiyoye cells**. (E-F)** P3HR1 cells stably expressing sh-RNA specific for E2F1 under doxycycline responsive promoter were **(E)** photographed using a Fluorescent Imager and subjected to **(F)** immunoblot analysis without or with doxycycline (-/+ DOX) treatment. **(G)** Immunoblot analysis of P3HR1 cells expressing either control (sgCon) or two E2F1 specific sgRNAs. **(H-I)** qRT-PCR of EBV intracellular or extracellular genome copy number from P3HR1 cells **(H)** without or with doxycycline (-/+ DOX) treatment and **(I)** in the presence of control or E2F1 sgRNAs. **(J-L)** P3HR1 cells stably expressing E2F1 sh-RNA in the absence and presence of doxycycline (-/+DOX) either left untreated or treated with TPA/sodium butyrate (NaBu). 72 h post treatment cells were subjected to **(J)** immunoblot, **(K)** qRT-PCR and **(L)** EBV intracellular or extracellular genome copy number analysis. The results are presented as the mean ± SD, n = 3 biological replicates. Statistical significance was determined by a two-sided Student’s t-test, *P < 0.05; **P < 0.01; ***P < 0.001; ns, not significant. Source data are available online for this figure.

In order to further confirm the phenomenon, P3HR1 cells were either knockdown (KD) for E2F1 using specific sh-RNA under doxycycline responsive promoter or knockout (KO) using two sets of sgRNAs (Fig 4E-G). The efficiency of KD/KO was validated using immunoblot analysis (Fig 4F-G). Intriguingly, either KD or KO of E2F1 markedly enhanced BZLF1 expression (Fig 4F-G) and virion production (Fig 4H-I) even without lytic cycle induction. To validate this notion further, P3HR1 cells stably expressed sh-RNA for E2F1 were either left untreated or subjected to viral lytic cycle induction by TPA/NaBu treatment in the absence or presence of doxycycline for 72 h (Fig 4J-L). While E2F1 KD or TPA/NaBu treatment alone enhanced BZLF1 transactivation and subsequent virion production, combination of both E2F1 KD and chemical lytic cycle inducer further enhanced the process (Fig 4J-L). In contrast, ectopic expression of E2F1 in HEK293T cells harbouring EBV bacmid robustly blocked EBV lytic gene expressions and mature virion production when lytic cycle was induced (Fig 5A-C). Overall, these results suggest that E2F1 specifically acts at the level of BZLF1 to govern EBV lytic cycle replication.

**Figure 5.**
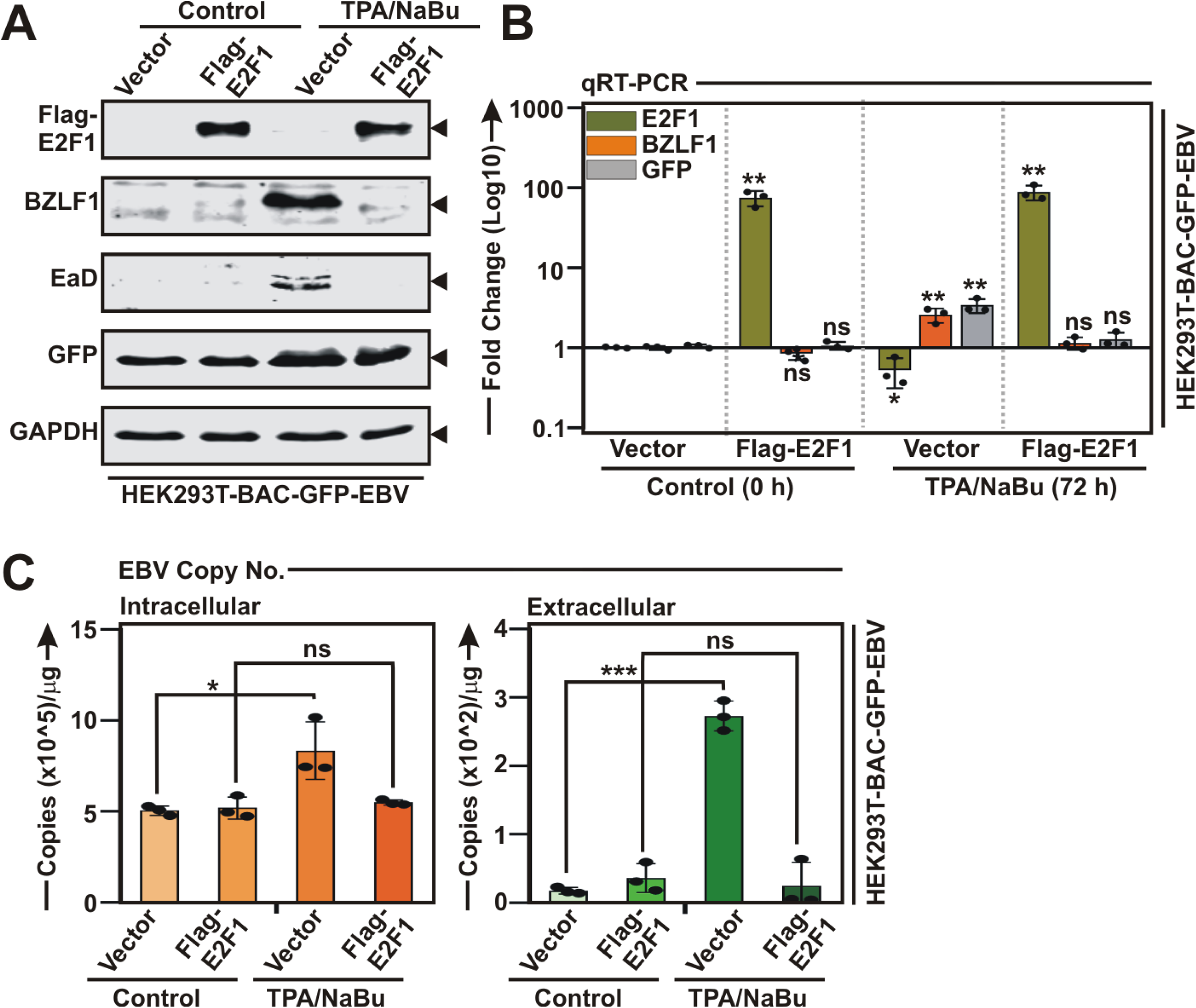
E2F1 overexpression retards EBV lytic cycle reactivation. **(A-C)** HEK293T-BAC-GFP-EBV cells transiently transfected either control vector or flag-tagged E2F1 expression plasmid, followed by TPA/sodium butyrate (NaBu) treatment for 72 h were subjected to **(A)** immunoblot, **(B)** qRT-PCR and **(C)** EBV intracellular or DNase-treated extracellular genome copy number analysis. The results are presented as the mean ± SD, n = 3 biological replicates. Statistical significance was determined by a two-sided Student’s t-test, *P < 0.05; **P < 0.01; ***P < 0.001; ns, not significant. Source data are available online for this figure.

### BZLF1-E2F1 cross-regulations are cell cycle and pRb independent

In contrast to most of the DNA viruses, herpesviruses including EBV induce cell cycle arrest at G1 phase during lytic cycle replication in both epithelial and B-cell backgrounds (Malik *et al*, 2024; Yockteng-Melgar *et al*, 2022). Although the underlying mechanisms are not well defined, it is suggested that this quasi-G1/S state expedites viral lytic replication. Notably, multiple EBV lytic cycle members including IE proteins BZLF1 and BRLF1, E protein BORF2 and L protein BGLF2 cause cell cycle arrest at the G1 to S phase transition (Rodriguez et al, 1999; Huang *et al*, 2012; Paladino *et al*, 2014; Yockteng-Melgar *et al*, 2022). This lytic replication-mediated cell cycle arrest prompted further investigation into its impact on E2F1 expression. Toward this objective, in addition to lytic cycle reactivation by TPA/NaBu, P3HR1 cells were additionally treated with mimosine/leucenol, a tyrosine analogue that specifically arrests dividing cells in the late G1 phase by inhibiting DNA replication initiation (Fig EV3A-C). In alignment with the previous findings, treatment with both mimosine and TPA/NaBu in P3HR1 cells exhibited characteristic G1/S cell cycle arrest (Fig EV3A-B). However, in contrast to EBV lytic cycle reactivation by TPA/NaBu treatment, mimosine-induced G1 arrest had no repressive effect on E2F1 expression levels (Fig EV3C).

While E2F1 is primarily associated with promoting cell cycle progression by transcriptional activation of genes essential for DNA replication and S-phase entry, it can also induce cell cycle arrest and apoptosis under various stress responses such as DNA damage signals. The retinoblastoma protein (pRb) negatively regulates both E2F1-driven cell cycle and apoptosis through interaction with a pRb-binding motif located at the edge of the C-terminal transactivation domain (residues 409-426). Nevertheless, ectopic expression of flag-tagged both WT (residues 1-437) and the mutant (residues 1-400) E2F1, lacking the pRb binding domain (PBD), in Jiyoye cells resulted in drastic depletion of basal BZLF1 of expressions (Fig EV3D-E), indicating that E2F1 mediated suppression of BZLF1 expression and subsequent viral lytic cycle reactivation is a pRb-independent phenomenon. Collectively, these results suggest that E2F1-BZLF1 cross-regulation is unaffected by cell-cycle arrest or its mediators.

### E2F1, but not E2F2, suppresses BZLF1 expression at the transcriptional level via its transactivation domain

The E2F TFs exerts their functions through specific DNA-binding and protein interacting domains (Fig 6A). Cell cycle activities of E2F isoforms are tightly regulated by pocket proteins (pRb, p107 and p130) dependent (E2F1-5) and independent (E2F6-8) manner. Apoptotic regulations of the prototype member E2F1 sets it apart from other E2Fs. Given both distinct and overlapping functions of E2F TFs, we next asked whether other E2F members can also suppress BZLF1 transactivation. Among the three activator family members E2F2 shared highest sequence homology (53%) with E2F1 (Fig 6A and Table EV3). Moreover, because in response to EBV lytic cycle induction, E2F1 and E2F2, but not E2F3, were significantly depleted at the transcriptional levels (Fig EV1G-H), we reasoned that E2F2 might have similar repressive activity on BZLF1 expression. However, in contrast to E2F1, ectopic E2F2 expression failed to inhibit BZLF1 expression as well as its promoter activity (Fig 6B-D). Computed structure model analysis demonstrated that although E2F1 and E2F2 share both significant primary sequence and three dimensional structure resemblance, exclusively within the DNA binding (DBD) and heterodimerization (DZD) domains, they display distinct features within the C-terminal transactivation domain (TAD) (Fig 6E). We therefore hypothesized that although E2F1 and E2F2 have similar DNA-binding abilities across the genome, owing to its specific TAD, E2F1 may display critical biological functions distinct from other E2Fs. Luciferase based promoter assay indeed revealed that WT (residues 1-437), but not, DTAD E2F1 (residues 1-358) lacking the TAD, failed to transcriptionally repress Zp activity (Fig 6F). To gain insights into E2F1 TAD negative regulations on BZLF1 expression, we swapped the respective TADs (residues 359-437) between E2F1 and E2F2 and generated constructs expressing chimeric proteins – E2F1-TAD2 (comprising E2F2 TAD) and E2F2-TAD1 (comprising E2F1 TAD). While contrary to WT E2F1, E2F1-TAD2 and WT E2F2 failed to transcriptionally repress Zp activity, E2F2 containing E2F1 specific TAD (E2F2-TAD1) significantly blocked Zp activity (Fig 6G). These data support a fundamental role of E2F1 TAD in transcriptional deactivation of BZLF1 during viral lytic replication.

**Figure 6.**
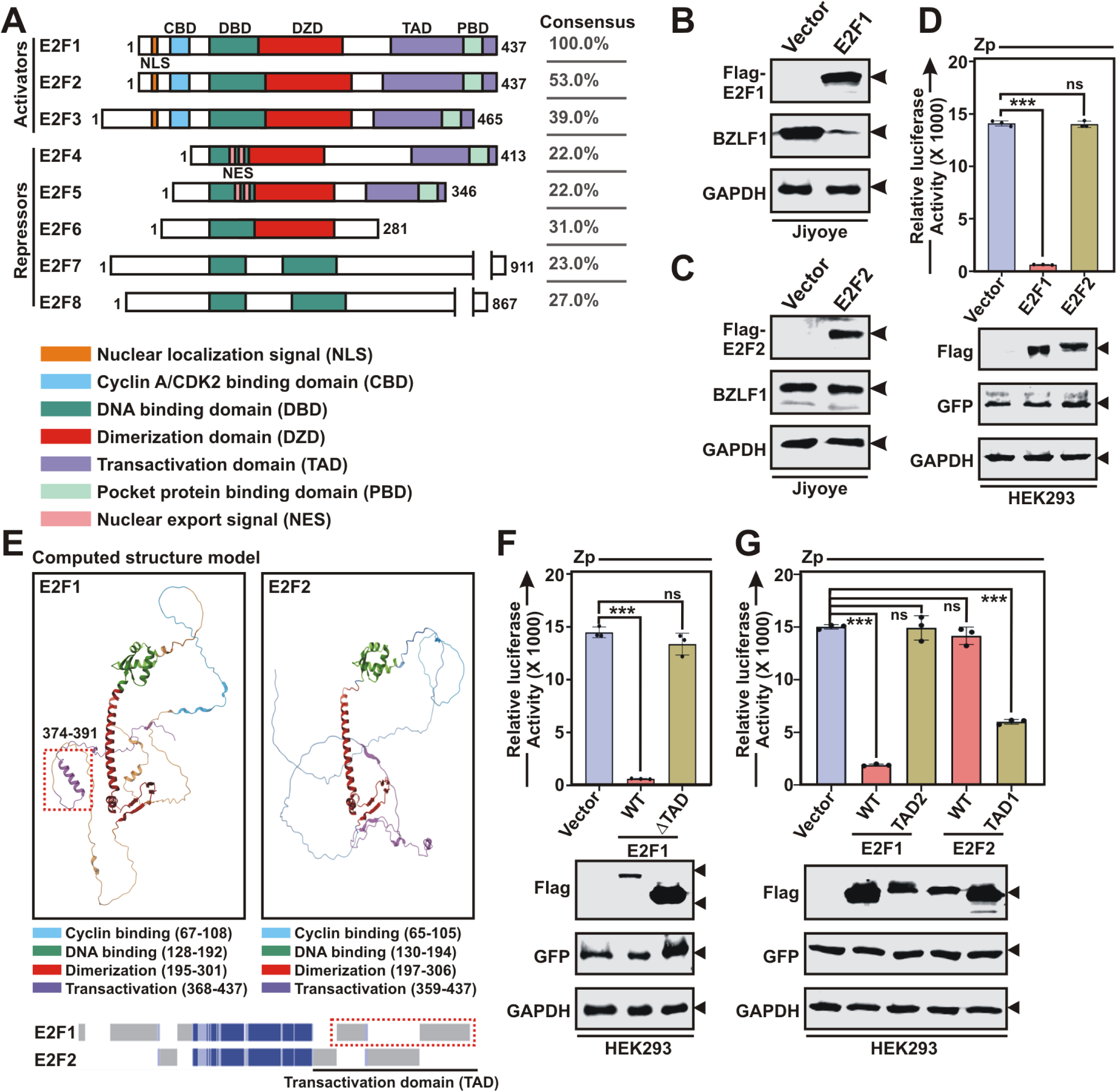
E2F1, but not E2F2, transcriptionally deactivates BZLF1 expression through its transactivation domain. **(A)** Schema showing known structural domains of all eight E2F isoforms (E2F1-8). Right panel indicates the % sequence similarities E2F1 and other E2F isoforms. **(B)** Immunoblot analysis of P3HR1 cells transiently transfected with either control vector or flag-tagged E2F1 expression plasmid. **(C)** Immunoblot analysis of P3HR1 cells transiently transfected with either control vector or flag-tagged E2F2 expression plasmid. **(D)** Luciferase reporter activity and the corresponding immunoblot analysis of the BZLF1 promoter (Zp) in the presence of control vector, or flag-tagged E2F1 or E2F2 expression plasmids. **(E)** Pairwise 3D structure alignment of E2F1 and E2F2. Bottom panel indicates sequence similarly (Blue) and dissimilarity (Grey) between E2F1 and E2F2 structure alignment. **(F)** Luciferase reporter activity and the corresponding immunoblot analysis of the Zp in the presence of empty vector, wild-type (WT) or transactivation domain deleted (ΔTAD) E2F1 expression plasmids. **(G)** Luciferase reporter activity and the corresponding immunoblot analysis of the Zp in the presence of empty vector, WT E2F1, E2F1 fused with E2F2-TAD domain (TAD2), WT E2F2 or E2F2 fused with E2F1-TAD domain (TAD1) expression plasmids. The results are presented as the mean ± SD, n = 3 biological replicates. Statistical significance was determined by a two-sided Student’s t-test, *P < 0.05; **P < 0.01; ***P < 0.001; ns, not significant. ND, not detected. Source data are available online for this figure.

### E2F1 expression restricts specific EBV latency program and lytic cycle reactivation

To further illustrate the effects of E2F1 on EBV gene expression, we performed RNA-Seq analysis of P3HR1 transcripts with or without E2F1 KD in the absence and presence of EBV lytic cycle induction following TPA/NaBu treatment (Fig EV4A and Dataset EV1). The transcriptomic data were first aligned with EBV type 2 genome. Most EBV lytic genes were transcriptionally activated by either E2F1 KD or TPA/NaBu treatment, combination of both E2F1 KD and the chemical inducer further amplified the effect (Fig EV4A and Dataset EV1). EBNA2 deleted P3HR1 usually displays atypical latency II consisting of both type-II (expressing LMP1 and LMP2A) and Wp-restricted (expressing EBNALP and EBNA3 genes - 3A/3B/3C) latency programs (Li *et al*, 2020). Among the latent genes, all three EBNA3 genes EBNA3A, EBNA3B and EBNA3C were transcriptionally activated upon E2F1 KD (Fig EV4A and Dataset EV1), indicating E2F1 depletion enforces a more Wp-restricted latency in P3HR1 cells. Additionally, Cp mediates transcription of EBNA genes in type-III latency cells (Murata *et al*, 2021). Consistent with this finding, luciferase based promoter assays demonstrated that while E2F1 expression significantly inhibited transcription from Wp and Cp, it facilitated transcription from Qp in a dose dependent manner (Fig EV2). Qp drives transcription of EBNA1 and LMP1/2 in latency IIa (Murata *et al*, 2021). Induction of EBV lytic and latent gene expressions in response to E2F1 depletion in P3HR1 cells was further validated by qRT-PCR analysis (Fig EV4B). While additional roles in EBV latency maintenance are possible, collectively these data suggest that E2F1 depletion favours a more Wp-restricted type-II or Cp-restricted type-III EBV latency programs and sensitizes lytic cycle reactivation. In agreement to this, RNA-Seq analysis of Mutu I and Mutu III cells (GSE136597), expressing Qp-restricted type-I latency and Cp-restricted type-III latency, respectively, revealed significant transcriptional repression of E2F1 in Mutu III cells (Fig EV4C).

### E2F1 acts prior to c-Myc for its positive regulation and thereby controls EBV latent-to-lytic switch

To gain insights into E2F1 transcriptional network involved in EBV lytic cycle reactivation, we further analysed the P3HR1 cell transcripts in response to E2F1 KD with or without EBV lytic cycle induction (Fig 7A and Dataset EV2). As expected, global transcriptomics analysis of both E2F1 KD and EBV lytic cycle reactivation demonstrated significant downregulation of genes involved in cell pathways featuring both G1/S and G2/M transitions of the cell cycle, DNA replication, B-cell activation, B-cell differentiation and B-cell receptor signalling (Fig 7A and Dataset EV2). Among the most downregulated genes by E2F1 depletion and lytic cycle induction, c-Myc appeared within top 10 genes (Fig 7B and Dataset EV2). This result raised the question of whether c-Myc TF, which is strongly expressed during EBV infection of naïve B-lymphocytes and thereby aids to latency establishment, could be a direct target of E2F1. Moreover, a recent genome-wide CRISPR/Cas9 screening identified c-Myc linked transcriptional network necessary for suppression of EBV lytic cycle replication (Guo *et al*, 2020a). While accumulating research indicated seemingly paradoxical transcriptional regulations of these two important molecules in various cell types, particularly in solid cancers (Coller *et al*, 2007; Fernandez *et al*, 2003; Ladu *et al*, 2008; Zhang *et al*, 2014), there are no reports available in the context of EBV associated B-cell lymphomas. DepMap analysis of 19 EBV positive B-cell lines demonstrated that E2F1 and c-Myc transcripts were significantly correlated (r=0.7067; p=0.001) (Fig 7C). In contrast, no significant correlation was found in EBV negative B-cell lymphoma lines as well as multiple epithelial cancer lines including lung, breast, prostate and skin carcinomas in DepMap cell lines portal (Fig S3A). Analysis of TCGA patients’ samples data of EBV negative diffuse large B-cell lymphoma (DLBCL) and corresponding solid cancers - lung adenocarcinoma (LUAD), breast invasive carcinoma (BRCA), prostate adenocarcinoma (PRAD) and skin cutaneous melanoma (SKCM) further validated this notion (Fig S3B).

**Figure 7.**
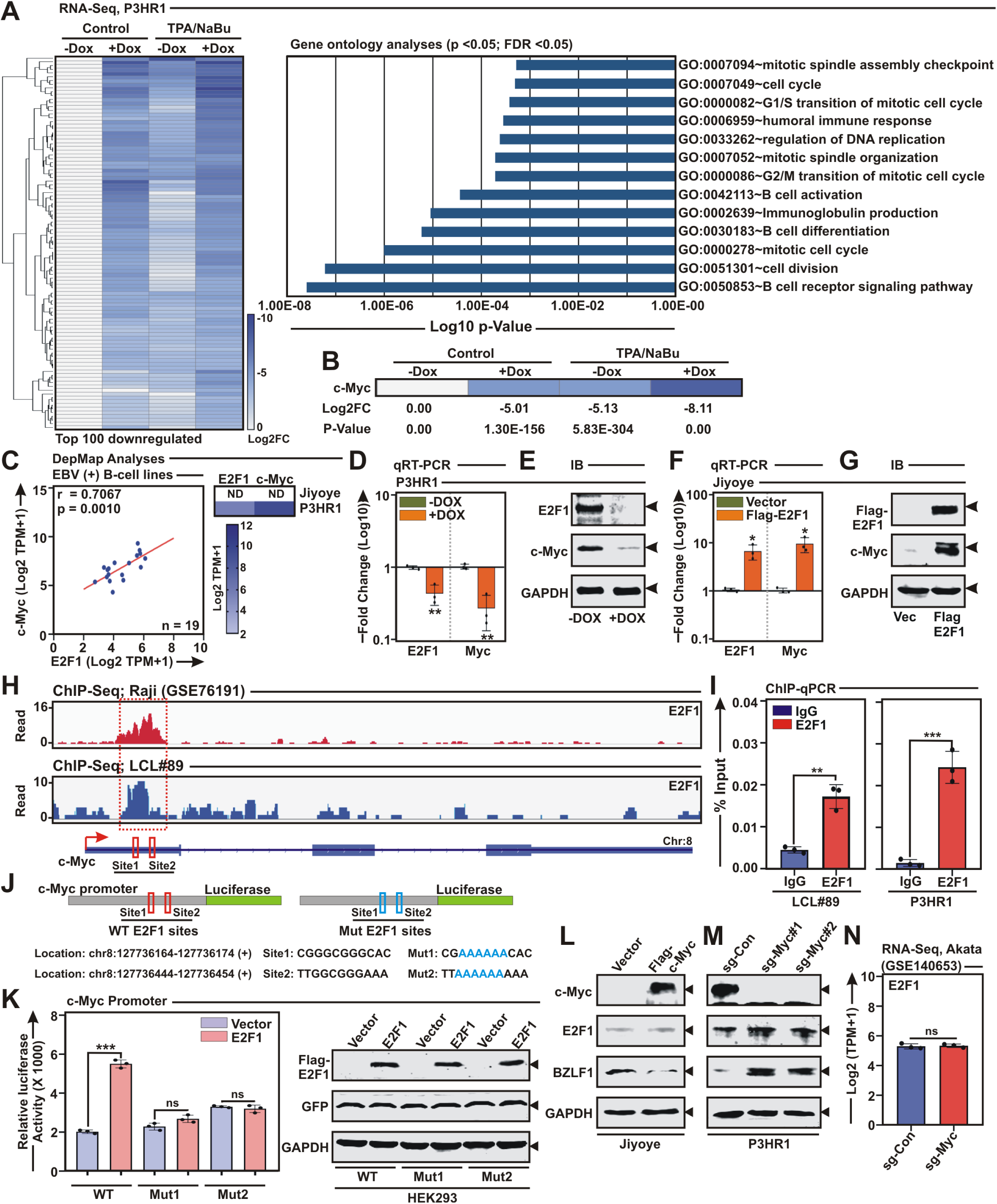
E2F1 positively regulates c-Myc expression. **(A)** Whole transcriptome analysis of P3HR1 stably expressing E2F1 sh-RNA in the absence and presence of doxycycline (-/+DOX) were either left untreated or treated with TPA/sodium butyrate (NaBu) for 72 h. Left panel indicates heatmap analysis of top 100 downregulated genes. Right panel bar diagram indicates most significantly affected pathways (p <0.05, FDR <0.05) based on top 100 downregulated genes in the RNA-Seq data of E2F1 knockdown and EBV lytic cycle reactivation. **(B)** Heat map visualization of c-Myc transcripts in the P3HR1 RNA-Seq data. **(C)** Two-sided unpaired Student’s t-test and two-sided Pearson’s correlation were employed to analyse the association between E2F1 and c-Myc transcripts in 19 EBV^+^ BL lines from DepMap portal. Right panel indicates heatmap representation of E2F1 and c-Myc expressions in P3HR1 and Jiyoye cells. **(D-E)** P3HR1 cells stably expressing E2F1 specific sh-RNA under doxycycline responsive promoter were subjected to **(D)** qRT-PCR and **(E)** immunoblot analysis without or with doxycycline (-/+ DOX) treatment. **(F-G)** Jiyoye cells transiently transfected with either control vector or flag-tagged E2F1 expression plasmid were subjected to **(F)** qRT-PCR and **(G)** immunoblot analysis. **(H)** EBV^+^ LCL#89 and Raji (GSE76191) ChIP-Seq tracks of E2F1 occupancy at c-Myc gene locus. **(I)** ChIP-qPCR analysis of E2F1 occupancy at c-Myc promoter region. **(J)** Schema showing two wild-type E2F1 binding sites (Red, Sites 1-2) and their corresponding mutations (Blue, Muts 1-2) on c-Myc promoter region for cloning into pGL3 luciferase reporter vector. **(K)** Luciferase reporter activity and the corresponding immunoblot analysis of the wild-type (WT) and mutant (Mut) c-Myc promoter in the absence and presence of flag-tagged E2F1 expression vector. **(L)** Immunoblot analysis of Jiyoye cells transiently transfected with either control vector or flag-tagged c-Myc expression plasmid. **(M)** Immunoblot analysis of P3HR1 cells expressing either control (sgCon) or two c-Myc specific sgRNAs. **(N)** Reanalysis of RNA-Seq data (GSE140653) of E2F1 transcripts in Akata cells expressing either control (sgCon) or c-Myc specific sgRNA. The results are presented as the mean ± SD, n = 3 biological replicates. Statistical significance was determined by a two-sided Student’s t-test, *P < 0.05; **P < 0.01; ***P < 0.001; ns, not significant. Source data are available online for this figure.

P3HR1 with elevated E2F1 expression displayed high level of c-Myc expression, while Jiyoye had insignificant expressions of both E2F1 and c-Myc (Fig 7C). We next asked whether E2F1 is directly responsible for c-Myc expression. Both qRT-PCR and immunoblot analysis of E2F1 depletion in P3HR1 and ectopic E2F1 expression in Jiyoye further confirmed the interdependence of E2F1 and c-Myc expressions in EBV positive B-cell lymphoma setting (Fig 7D-G). To gains insights into the underlying mechanism governing E2F1-directed c-Myc transcription, we analysed/re-analyzed E2F1 ChIP-Seq data-sets in both LCLs and BL background (GSE76191) (Fig 7H). ChIP-Seq profiles indicated strong E2F1 enrichment in the first exon of the c-Myc gene locus (Fig 7H). E2F1 ChIP-Seq signals on c-Myc promoter/enhancer were further validated by ChIP-qPCR analysis in both LCLs and P3HR1 cells (Fig 7I). To further assess the E2F1-mediated transcriptional regulation of c-Myc, we performed luciferase reporter assay using the c-Myc exonic region spanning from 127736027 to 127736461 in chromosome 8 containing two putative E2F1 binding motifs (Fig 7J-K). A concentration dependent increase in the c-Myc promoter driven luciferase activity was observed in HEK293 cells transfected with E2F1 expression vector (Fig EV5A). While WT E2F1 significantly increased luciferase activity, mutant E2F1 with deleted TAD (E2F1 ΔTAD) failed to transactivate the c-Myc promoter/enhancer region (Fig EV5B). Luciferase reporter assay with mutations in both E2F1-recognition sites within c-Myc locus further established E2F1 mediated positive transcriptional regulation of c-Myc (Fig 7J-K). Together, these findings indicated that the E2F1 acts as a direct transcriptional activator of c-Myc.

Since c-Myc and E2F1 have been shown to activate each other’s transcription, we next investigated the impact of c-Myc on E2F1 transcription in EBV positive cells. In contrast to the effect of ectopic E2F1 expression on increased c-Myc expression in both Jiyoye and HEK293T cells harbouring EBV bacmid (Figs 7G and EV5C), c-Myc expression failed to elevate endogenous E2F1 expression in these cells (Figs 7L and EV5D). To further validate this data, two sgRNAs were designed to target c-Myc and subsequently checked their effect on E2F1 expression in P3HR1 cells (Fig 7M). In agreement with the ectopic expression settings, c-Myc depletion also had no effect on E2F1 expression (Fig 7M). RNA-Seq analysis (GSE140653) of EBV positive Akata transcripts following expression of control or sgRNA targeting c-Myc further validated this notion (Fig 7N). Additionally, analysis of publicly available two ChIP-Seq datasets (GSE30399 and GSE36354) for c-Myc in LCLs revealed no distinct peaks in E2F1 gene locus (Fig EV5E). In concordance with these data, no change in the luciferase activity was noted in HEK293 cells transfected with WT E2F1 promoter in the absence and presence of increasing doses of c-Myc expressing construct (Fig EV5F), suggesting that E2F1 acts prior to c-Myc for its transcriptional activation.

Consistent with prior reports of c-Myc depletion mediated EBV lytic cycle reactivation in Akata cells (Guo *et al*, 2020a), c-Myc KO by both sgRNAs significantly elevated BZLF1 expression in P3HR1 cells (Fig 7M). Notably, although c-Myc was proposed to act at the level of BZLF1 through chromosome looping to OriLyt region to control EBV lytic replication (Guo *et al*, 2020a), no direct binding of c-Myc was established in the BZLF1 promoter/Zp region. After inspection of EBV Zp using JASPAR database, in contrast to three E2F1 binding sites, one putative c-Myc binding site (CCACCTG/C) was identified (Fig 8A). While significant decrease in luciferase activities were observed in HEK293 cells transfected with Zp in the presence of either E2F1 or c-Myc expression construct in a dose dependent manner, as opposed to c-Myc, E2F1 demonstrated considerably higher suppressive activity of Zp (Fig 8B). The c-Myc-mediated transcriptional repression of Zp activity was further confirmed by mutational analysis of the single c-Myc binding site (Fig 8C). Additionally, no synergistic effect between c-Myc and E2F1 mediated transcriptional suppression of Zp activity was observed (Fig 8B-C). Depletion of c-Myc expression during lytic cycle reactivation (Figs 7B and S4A and Dataset EV2) combined with BZLF1 mediated E2F1 transcriptional repression (Fig 2A-J) prompted us to further investigate BZLF1’s effect on c-Myc transcription (Fig S4B-C). However, ChIP-Seq profile (E-MTAB-7788) and luciferase reporter analysis demonstrated neither BZLF1 occupies in the c-Myc promoter region nor it influence c-Myc promoter activity (Fig S4B-C). In sum, these results are consistent with a model in which E2F1 transcriptionally activates c-Myc and they independently repress BZLF1 expressions during EBV latent infection of naïve B-lymphocytes, whereas in response to periodic lytic cycle reactivation E2F1 and BZLF1 are present in a loop and negatively modulate each other’s expression (Fig 8D). BZLF1-driven restricted E2F1 expression enables in lowering the c-Myc levels, thereby supporting the maintenance of EBV lytic replication (Fig 8D).

**Figure 8.**
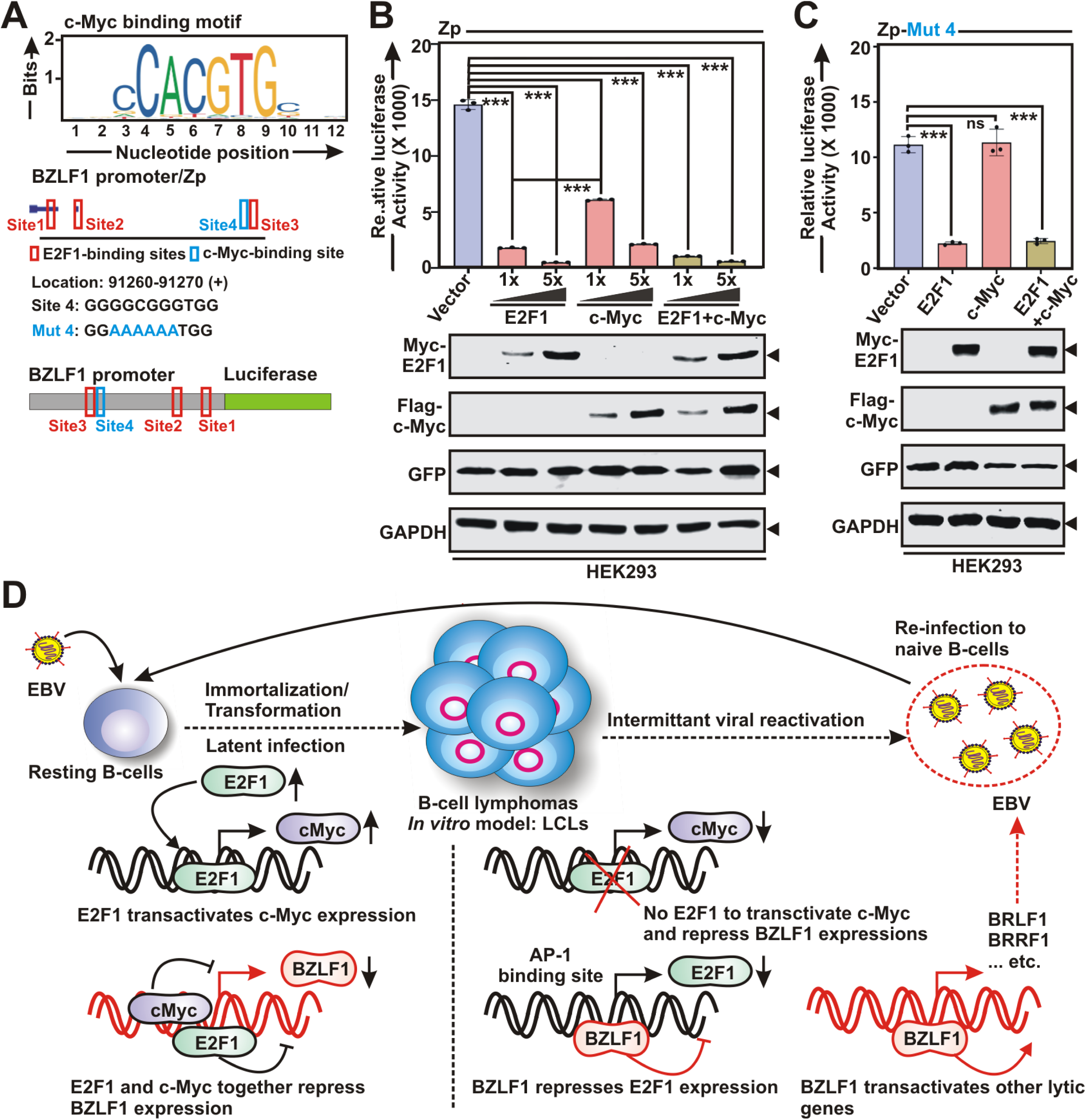
Together with c-Myc, E2F1 transcriptionally repress BZLF1 expression and EBV lytic cycle reactivation. **(A)** E2F1 (red, sites 1-3) and c-Myc (blue, site 4) binding motifs obtained from JASPAR database in BZLF1 promoter (Zp) region. Schema showing single wild-type c-Myc binding site (Site 4) and the corresponding mutation (Mut 4) on Zp for cloning into pGL3 luciferase reporter vector. **(B-C)** Luciferase reporter activity and the corresponding immunoblot analysis of the **(B)** wild-type and **(C)** mutant Zp in the absence and presence of E2F1 and c-Myc expression plasmids either in increasing concentrations or in combination. **(D)** Schematic model of the roles of E2F1 in repressing EBV lytic cycle reactivation. During EBV latency, E2F1 promotes c-Myc expression, together which bind to the BZLF1 promoter and repress its leaky expression that impedes subsequent expression of other lytic genes. Upon EBV lytic reactivation, BZLF1, on the contrary, transcriptionally repress E2F1 and thereby c-Myc expression, highlighting a unidirectional regulatory hierarchy. BZLF1 mediated E2F1 depletion deploys an effective lytic cycle environment that facilitates a cascade of EBV lytic gene expression. The results are presented as the mean ± SD, n = 3 biological replicates. Statistical significance was determined by a two-sided Student’s t-test, *P < 0.05; **P < 0.01; ***P < 0.001; ns, not significant. Source data are available online for this figure.

## Discussion

In human, EBV establishes a life-long infection rendering a biphasic viral life cycle – latent period and lytic replication. While latent phase allows EBV to evade the host’s immune surveillance and develop oncogenic phenotypes, periodic lytic reactivation is essential for viral progeny production and horizontal transmission (Shannon-Lowe *et al*, 2017; Yap *et al*, 2022). Moreover, accumulating evidence suggests that EBV lytic proteins contribute to cancer progression (Dorothea *et al*, 2023; Hong *et al*, 2005). Despite its critical role in EBV pathogenesis, the molecular mechanisms governing the latent-to-lytic switch remain poorly understood. For example, studies suggest that terminal differentiation of memory B-lymphocytes into plasma cells initiates spontaneous EBV replicative cycle (Laichalk & Thorley-Lawson, 2005). Moreover, recently, it has been demonstrated that c-Myc expression levels controls EBV latent-to-lytic switch by altering viral three-dimensional genome architecture (Guo *et al*, 2020a). Our study provides critical insights into the regulatory dynamics between E2F1 and c-Myc transcription factors, elucidating their roles in EBV lytic cycle reactivation. By dissecting the mechanisms through which E2F1 suppresses EBV lytic replication and interacts with cellular and viral gene networks, our findings provide deeper insights into EBV pathogenesis and inform potential therapeutic interventions.

E2F1 regulates both cell cycle and apoptosis, and its activity is often deregulated in human tumours (Kumari *et al*, 2015; Shats *et al*, 2017). Oncogenic viruses employ a common strategy for E2F1 transactivation, which enables continuous cell proliferation. For example, simian SV40 virus encoded LT, adenovirus encoded E1A and human papilloma virus (HPV) encoded E7 disrupt the interaction between E2F1 and pRb to facilitate G1 to S phase transition of the cell cycle (DeCaprio *et al*, 1988; Mal *et al*, 1996; Hwang *et al*, 2002). However, in case of EBV, the scenario is much complex and inadequately represented. Alike various viral oncoproteins, EBV encoded EBNA3C interacts with pRb and expedites its ubiquitin mediated degradation (Knight *et al*, 2005), raising the possibility of manipulating E2F1 mediated oncogenic activities. Instead, EBNA3C inhibits E2F1 mediated apoptosis under DNA-damage signals and recruits E2F6 for its transcriptional repression (Saha *et al*, 2012; Pei *et al*, 2016). Moreover, EBNA1, the only viral oncoprotein expressed in LCLs, all varieties of EBV-associates BL and other solid tumours such as nasopharyngeal carcinoma (NPC) and gastric carcinoma (EBVaGC), activates E2F1 expression at post-transcriptional level via PI3Kδ-dependent mRNA translation stress (Gnanasundram *et al*, 2017). Reanalysis of our previous lab and publicly available genome-wide transcriptome data revealed E2F1 activation during EBV infection of primary B-lymphocytes and BL cells. DNA damage responses elicit cell cycle arrest and facilitate DNA repair, ensuring overall genomic integrity. However, owing to incessant cell proliferation as well as faulty DNA repair pathways, cancer cells often exhibit ‘replication stress’, which leads to genomic instability (Gaillard *et al*, 2015). E2F1 localizes at these DNA break sites and subsequently repairs them by recruiting homologous recombination factors (Choi *et al*, 2019). It has been suggested that the ‘replication stress’ upon EBV infection represents a major barrier for naïve B-cell transformation (Nikitin *et al*, 2010). EBNA3C can impede this EBV induced DNA damage response and thereby enabling B-cell growth transformation (Nikitin *et al*, 2010). Although the underlying mechanism is still incomplete, it is conceivable that during latent infection EBV critically modulates E2F1 levels to mitigate DNA damage response and promote efficient B-cell transformation.

In contrast to latent infection, our data demonstrated that EBV lytic cycle replication transcriptionally repressed E2F1 expression. The lytic phase is initiated by leaky expression of the IE gene, BZLF1, a key driver of lytic cycle replication, which subsequently transcriptionally activates a cascade of 30 lytic genes (Bernaudat *et al*, 2022; Guo *et al*, 2020a). Interestingly, BZLF1 demonstrates a propensity to bind and transactivate CpG methylated viral promoters to mitigate epigenetic silencing during latent-to-lytic switch (Bhende *et al*, 2004; Bergbauer *et al*, 2010). As opposed to BZLF1, the other IE protein BRLF1 supports lytic replication by promoting lytic gene expression from hypomethylated viral promoters (Wille *et al*, 2013). Our findings establish E2F1 as a fundamental regulator of EBV latency through intervening lytic replication by transcriptionally repressing BZLF1 expression in both B-lymphocyte and epithelial cell background. In agreement to this, overexpression of E2F1 along with c-Myc inhibit BZLF1 mediated transactivation (Lin *et al*, 2004). The transcriptional repression mediates through a negative regulatory element located at the N-terminal transactivation domain of BZLF1 (Lin *et al*, 2004). Since treatments with DNA-methylation inhibitor and HDAC inhibitor promote BZLF1 expression and subsequent lytic replication (Li *et al*, 2012; Murata *et al*, 2012), it is therefore conceivable that epigenetic regulation plays critical roles in lytic cycle reactivation. Recently, genome-wide CRISPR/Cas9 screening identified UHRF1, an E3 ubiquitin ligase, and its DNA methyltransferase partner DNMT1 as critical determinants for EBV latency programs (Guo *et al*, 2020b). Towards stably maintaining DNA methylation, DNMT1 requires UHRF1 (Nishiyama *et al*, 2020). UHRF1 depletion transforms latency I to latency III by de-repressing EBNA and LMP expressions (Guo *et al*, 2020b). UHRF1 silencing also robustly increases BZLF1 expression in EBV positive cells (Guo *et al*, 2020b). Importantly, both UHRF1 and DNMT1 are the transcriptional targets of E2F1 (Benavente *et al*, 2014; McCabe *et al*, 2005; Li *et al*, 2019), where activated MEK/ERK pathway functions as a driving force (Li *et al*, 2019). We speculate that in addition to direct transcriptional regulation, E2F1 may also be involved in epigenetic regulation of BZLF1 repression through adjusting UHRF1 expression level. In agreement to these, reanalysis of RNA-Seq data (GSE237484, GSE136597) demonstrated that several E2F1 targeted genes including UHRF1 expressions were significantly downregulated in latency III EBV positive B-cells as well as in response to EBV lytic cycle reactivation. This also raises the possibility of exploiting UHRF1 inhibitors (Chang *et al*, 2021) as a potential therapeutic strategy for EBV-associated B-cell malignancies by inhibiting E2F1 downstream activities and thereby inducing EBV lytic replication.

Our study supports a model in which E2F1 and BZLF1 forms a negative transcriptional feedback loop, ensuring a tightly regulated transition between EBV latency and lytic replication. While during latent phase E2F1 transcriptional network is maintained above a threshold level to suppress BZLF1, we speculate that leaky BZLF1 expression owing to spontaneous B-cell differentiation into plasmocytes (Reusch *et al*, 2014) is enough to counteract E2F1 activities, thus orchestrating efficient viral lytic replication. Interestingly, BLIMP1, a master regulator of plasma cell development, positively regulates BZLF1 expression, also transcriptionally represses mature B-cell gene expression program including E2F1 (Shaffer *et al*, 2002). Likewise, another essential transcription factor for plasma cell differentiation XBP-1 stimulates BZLF1 expression (Bhende *et al*, 2007), may also repress E2F1 activity by an incompletely characterized mechanism (Obiedat *et al*, 2019). These findings are consistent with our model implicating the inverse correlation of E2F1 and BZLF1 as a key determinant of contradictory EBV-infected B-cell states.

The regulatory axis involving E2F1 and c-Myc adds another layer of complexity to EBV latency maintenance. Our results demonstrate that during latency in addition to suppress BZLF1 expression, E2F1 positively regulates c-Myc level. However, c-Myc does not reciprocally regulate E2F1, underscoring a unidirectional regulatory hierarchy. This interplay highlights the role of c-Myc as a co-regulator of EBV latency, acting in concert with E2F1 to maintain suppression of the lytic cycle. Accumulating evidence strongly implicates c-Myc as one of the key factors in transforming naïve B-lymphocytes as well as sustaining viral latency by suppressing BZLF1 transcription (Guo *et al*, 2020a; Lin *et al*, 2004; Wang *et al*, 2019). In conjunction with previous data, our results also demonstrate that c-Myc depletion enhances BZLF1 expression and thereby accelerates EBV lytic replication. While several reports indicated cross-regulation between E2F1 and c-Myc (Coller *et al*, 2007; Ladu *et al*, 2008; Zhang *et al*, 2014), to our knowledge, has not previously been implicated in EBV associated B-cell malignancies. Our study provides compelling rationale for repressing EBV lytic cycle replication, where E2F1 forms a feedforward loop for transcriptional activation of c-Myc, together which limit BZLF1 expression and thereby lytic reactivation. Our data supports a lytic reactivation model, where BZLF1’s ability to repress E2F1 and thereby c-Myc transcription ensures a sustained lytic replication phase, while it fails to directly modulate c-Myc expression.

While our study provides a robust framework for understanding E2F1’s role in EBV reactivation, several questions remain unanswered. The mechanisms underlying cell type-specific differences in E2F1 activity and its interplay with other transcription factors during EBV infection warrant further investigation. Our study was exclusively focussed on E2F1 downstream modulator c-Myc, as evident from the top 100 hits of E2F1 knockdown P3HR1 cells. However, there are strong possibilities that additional hits may also function as potential repressors of EBV lytic reactivation. For example, cellular kinase and cyclin B partner CDK1 was identified as an interacting partner of EBV lytic cycle kinase BGLF4 (Zhu *et al*, 2009). It has been suggested that while BGLF4 mediated kinase activity is essential for an effective lytic replication, it significantly interferes EBNA1-directed latent replication (Zhu *et al*, 2009). Another EBV lytic protein BDLF4 phosphorylation by S-phase cyclin complexes – cyclin A/CDK2 and cyclin E/CDK2 but not by cyclin B/CDK1 is important to initiate viral late gene expression (Sato *et al*, 2019). We speculate that cyclin B/CDK1 complex is important for EBV latency maintenance and it will therefore be of interest to examine CDK1 depletion in viral lytic cycle reactivation. Additionally, understanding how chromatin remodelling influences E2F1-mediated transcriptional regulation could provide deeper insights into its role in EBV latency maintenance and reactivation into lytic cycle replication. An objective of future studies will be to delineate how both positive and negative upstream regulators of E2F1 control EBV latent-to-lytic switch program.

Our findings accentuate the therapeutic potential of targeting E2F1 and its regulatory circuits to modulate EBV lytic replication and treat associated B-cell malignancies. In sum, our study elucidates the intricate transcription network of E2F1/c-Myc axis, which governs the latent-to-lytic switch in EBV positive cells. As similar to c-Myc (Guo *et al*, 2020a), our results also demonstrated that E2F1 depletion significantly enhances virion production, even in the absence of chemical inducers. This finding aligns with the therapeutic concept of ‘lytic induction therapy’, which aims to transition latent EBV-infected cells into the lytic phase, rendering them susceptible to antiviral treatments (Tan *et al*, 2023; Yiu *et al*, 2020). Thus, the ability to modulate E2F1 transcription network with minimal cytotoxicity holds promise for improving the clinical management of EBV-associated neoplasms.

## Materials and Methods

### Cell lines and reagents

HEK293, Lenti-X 293T cells were cultured in Dulbecco’s modified Eagle’s medium (DMEM) (Gibco) supplemented with 10% FBS (Gibco) and 1% Penicillin-Streptomycin Solution (Gibco). HEK293T-BAC-GFP-EBV cells (Halder *et al*, 2009) were maintained in complete DMEM containing 1 µg/ml puromycin (Merck). EBV positive BL lines - P3HR1, Jiyoye, Namalwa and EB3 were obtained from NCCS cell repository, Pune, India (https://www.nccs.res.in/cellrepository). EBV transformed lymphoblastoid cells - LCL#1 and LCL#89 (Malik *et al*, 2024) and BL lines were maintained in RPMI 1640 (Gibco) supplemented with 10% FBS and 1% Penicillin-Streptomycin Solution. All cells were cultured at 37^0^C in a humidified environment with 5% CO_2._ All cell lines were routinely tested for *Mycoplasma* by LookOut Mycoplasma qPCR Detection Kit (Merck).

### Bioinformatic analysis of sequencing data

All raw sequencing reads from high throughput sequencing pipelines were first checked using FastQC (http://www.bioinformatics.babraham.ac.uk) and confirmed with no significant quality issues.

#### RNA-Seq data analysis

The raw Fastq files of transcriptome data of EBV infected PBMCs 0-4 DPI (GSE235941), EBV infected naïve B-lymphocytes 0-28 DPI (GSE125974), EBV lytic cycle reactivation from LCLs (GSE237484), Mutu I and Mutu III (GSE136597) and Akata c-Myc KO (GSE140653) were downloaded and extracted with unique SRR accession in Galaxy webserver (https://usegalaxy.org/) and complete analysis was performed using appropriate tools offered by the webserver. Data were visualized with either GrapPad Prism v8 or Microsoft Excel.

#### ChIP-Seq data analysis

The raw Fastq files of different ChIP-Seq data (BZLF1: E-MTAB-7788; E2F1: GSE76191; EBNA1: GSE73887; EBNA2: GSE29498; EBNALP: GSE49338; EBAN3A: GSE88729; EBNA3B: GSE88729; EBNA3C: GSE88729; RelA/p65: GSE55105; c-Rel: GSE55105; RelB: GSE55105; p50: GSE55105; p52: GSE55105; c-Myc: GSE30399 and GSE36354) were extracted from SRA files and aligned against human reference genome (Homo sapiens.GRCh37) using default parameters. BAM files were further analysed for peak calling using MACS2. Analysed files were visualized on the Integrative Genomics Viewer (https://igv.org/).

#### DepMap data analysis

To perform correlation study between E2F1 and c-Myc expressions across different EBV positive and negative B-cell lymphoma cell lines along with multiple solid cancers – lung, breast, prostate and skin carcinomas, gene expression data were extracted from DepMap Expression Public 23Q4 dataset (https://depmap.org/portal/). Correlation data was visualized using GrapPad Prism v8.

#### TCGA data analysis

To check the correlation between E2F1 and c-Myc expressions in diffuse large B-cell lymphoma (DLBCL), lung adenocarcinoma (LUAD), breast invasive carcinoma (BRCA), prostate adenocarcinoma (PRAD) and skin cutaneous melanoma (SKCM) patients’ samples UCSC Xena browser (https://xenabrowser.net/) was utilized. Extracted gene expression data were further analysed using GrapPad Prism v8 data.

#### GTEx data analysis

The Genotype-Tissue Expression (GTEx) Portal (https://www.gtexportal.org/home/) was utilized to check the expression pattern of all E2F isoforms between EBV transformed B-cells (LCLs) and whole blood samples (PBLs).

### EBV infection of PBMCs

HEK293T-BAC-GFP-EBV cells were induced for EBV lytic cycle reactivation with 20 ng/ml 12-O-Tetradecanoylphorbol-13-acetate (TPA; Merck) and 3 mM Sodium butyrate (NaBu; Merck) for 5 days. EBV particle in the culture supernatant was concentrated by ultracentrifugation at 27,000 rpm for 2 h at 4^0^C and subsequently re-suspended in 500 µl RPMI 1640 without any supplementation. Virus stock was stored at −80^0^C for further use. ∼1.0 x 10^7^ PBMCs (HiMedia) from two individual donors in complete RPMI 1640 were incubated with EBV (MOI: ∼10). 24 h post-infection, cells were centrifuged, re-suspended in fresh RPMI 1640 and continued growing at 37^0^C for 28 days. Cells were harvested at the indicated time points (0-28 DPI; days post-infection) for qRT-PCR analysis. EBV lytic cycle reactivation in HEK293T-BAC-GFP-EBV cells and latent infection in PBMCs were monitored by checking green fluorescence using a ZOE Fluorescent Cell Imager (BIO-RAD).

### Quantitative real-time (qRT-PCR) analysis

For qRT-PCR analysis, ∼1.0 x 10^7^ cells from each experimental settings were harvested for RNA isolation using PureZOL RNA Isolation reagent (BIO-RAD) following the manufacturer protocol. ∼1 µg RNA was subjected to reverse transcription using iScript cDNA synthesis kit (BIO-RAD) according to the manufacturer’s protocol. Both quality and quantity of nucleic acids were checked in a Synergy H1 Multimode Microplate Reader (BioTek). qRT-PCR analysis was conducted using iTaq Supermix (BIO-RAD) on a CFX Connect real-time PCR detection system (BIO-RAD). Unless specified otherwise, each reaction was replicated thrice and relative transcript levels were quantified using the 2^−ΔΔCT^ method and normalized with GAPDH or B2M internal control. All samples were run in technical triplicates and at least two independent experiments were performed.

### Immunoblot analysis

For immunoblot analysis, ∼1.0 x 10^7^ cells were lysed in 500 µl RIPA buffer (Thermo Fisher Scientific) combined with 1x protease inhibitor cocktail (Abcam) by occasional vortexing for 15 s with 5 min interval. After estimation of total protein concentrations by Bradford reagent (BIO-RAD), protein samples were boiled with 2x laemmli buffer (BIO-RAD) at 95^0^C for 10 min. Equal amount of samples were resolved by SDS-PAGE, transferred to a nitrocellulose membrane (BIO-RAD) and blocked with 5% milk in 1x TBS. After washing with 1x TBST, the membranes were incubated with specific primary antibodies overnight at 4^0^C. Following day, the membranes were washed with 1x TBST and incubated with appropriate infrared-tagged/DyLight secondary antibodies (Thermo Fisher Scientific) for 1 h at room temperature. After washing with 1x TBST, image analysis and quantification of protein bands were achieved using the Odyssey Infrared Imaging System (LiCor Inc.)

### EBV lytic cycle reactivation

For induction of EBV lytic cycle replication, ∼1.0 x 10^7^ HEK293T cells stably transfected GFP-tagged EBV-BACmid (HEK293T-BAC-GFP-EBV), EBV positive BL lines (P3HR1, Jiyoye, EB-3) and *in vitro* EBV transformed LCLs (LCL#89) were maintained in complete media containing either combination of 20 ng/ml TPA (Merck) and 3 mM NaBu (Merck) or treated with 1 μM MG132 (Abcam) or 10 µg/ml Goat Anti-Human IgG (Abcam) as indicated. 24-72 h post-treatment the lytic reactivation was validated by either immunoblotting or qRT-PCR analysis of EBV lytic cycle transactivator BZLF1 as well as quantification of EBV genome copy number.

### ChIP-qPCR analysis

ChIP-qPCR was performed as previously described (Malik *et al*, 2024). Briefly, after crosslinking and subsequent de-crosslinking ∼2.0 x 10^7^ cells were harvested, washed with ice-cold 1x PBS and suspended in lysis buffer (50 mM Tris-HCl pH 8.1, 10 mM EDTA, 1% SDS and 1 × protease inhibitor cocktail). Chromatin was sonicated with a Diagenode Bioruptor Plus sonicator (Diagenode Inc.) to attain DNA fragments of ∼200 to 400 bp as confirmed by agarose electrophoresis. 10% of the sonicated chromatin was collected and used as input material, while the remaining sheared chromatin was further diluted to immunoprecipitation (IP) dilution buffer (16.7 mM Tris-HCl pH 8.1, 1.2 mM EDTA, 167 mM NaCl, 1.1% Triton X-100, 0.01% SDS along with 1 × protease inhibitor cocktail), followed by immunoprecipitation with 5 μg appropriate antibodies (anti-E2F1 or anti-BZLF1) or corresponding mouse IgG control using magnetic protein A/G beads (BIO-RAD). After sequential washing steps with ‘low-salt wash buffer’ (20 mM Tris-HCl, pH 8.1, 2 mM EDTA, 150 mM NaCl, 1% Triton X-100, 1% SDS), ‘high-salt wash buffer’ (20 mM Tris-HCl, pH 8.1, 2 mM EDTA, 500 mM NaCl, 1% Triton X-100, 1% SDS)), ‘LiCl wash buffer’ (10 mM Tris-HCl, pH 8.1, 1 mM EDTA, 0.25 M LiCl, 1% NP-40, 1% deoxycholate acid) and ‘TE buffer’ (10 mM Tris-HCl, pH 8.1, 1 mM EDTA) the protein-DNA complexes were eluted using ‘elution buffer’ (100 mM NaHCO3, 1% SDS). Following reverse cross-linking using proteinase K treatment, the ChIP-ed DNA was purified using the QIAquick PCR purification kit (QIAGEN) and subjected for qPCR analysis. Data were analyzed by the ΔΔCT method relative to input DNA and normalized to the IgG control.

### Transfection

For transient transfection assays, ∼2.0 x 10^7^ Jiyoye, or ∼1.0 x 10^7^ HEK293 or HEK293T-BAC-GFP-EBV cells were harvested and re-suspended in 450 μL Opti-MEM (Gibco), mixed with appropriate plasmids in Gene Pulser/MicroPulser Electroporation Cuvettes (BIO-RAD) followed by electroporation using Gene Pulser II electroporator (BIO-RAD). For Jiyoye, electroporation pulses were set at 240 V and 960 μF, while for HEK293 and HEK293T-BAC-GFP-EBV cells pulses were set at 210 V 975 μF. Unless and otherwise stated, 36-48 h post-transfection cells were harvested and subjected for expression analysis. For promoter assays, HEK293 cells were transfected with appropriate plasmids using JetPrime (Polyplus Transfection Inc.) according to manufacturer’s protocol. For lentivirus production, Lenti-X 293T cells were transfected with appropriate plasmids using calcium phosphate transfection kit (Thermo Fisher Scientific).

### Co-immunoprecipitation (Co-IP)

Co-IP was performed as described previously (Gain *et al*, 2020). Briefly, ∼15 x 10^6^ HEK293 cells transiently transfected with myc-tagged BZLF1 with or without flag-tagged E2F1 expression plasmids were harvested, washed with 1 x PBS and subsequently lysed with 500 µl RIPA buffer supplemented with protease inhibitor cocktail. After saving 5% of the lysate as input, remaining lysate was subjected to preclear with Protein-A/G magnetic beads (BIO-RAD) for 1h at 4^0^C. Protein of interest was captured by rotating precleared lysate with 1 μg of anti-flag mouse monoclonal antibody (Merck) overnight at 4^0^C. Following day, Immuno-complexes were captured by Protein-A/G magnetic beads, washed with RIPA buffeter for three times and boiled with 2 x laemmli buffer for 5 min. Input lysates and IP complexes were then fractionated by SDS-PAGE and subjected to immunoblot analysis as mentioned above. The data was analysed and viewed on an Odyssey CLx Imaging System.

### Construction of plasmids and Site-Directed Mutagenesis

Cellular E2F1 and c-Myc wild type promoters along with all the EBV promoters - Zp, Cp, Qp, Wp, LMP1p and LMP2p were constructed in pGL3 Basic Vector (Addgene #212936) by conventional PCR, restriction digestion with appropriate enzymes followed by ligation. Substitution based mutated primers were designed using NEBaseChanger v2.5.2 tool (https://nebasechanger.neb.com/). Primers were obtained from Integrated DNA Technologies and mutated plasmids were constructed using Q5 Site-Directed Mutagenesis Kit (New England Biolabs) according to manufacturer’s protocol. Transactivation domains (TAD) of E2F1 and E2F2 were swapped using assembly based PCR with appropriate primers, followed by restriction digestion and ligation in pCDNA3.1 based vector (pA3F with 3x Flag). All the constructs were further verified by Sanger dideoxy based DNA sequencing (Eurofins Genomics India Pvt. Ltd., India).

### Luciferase-based promoter assay

Dual-Glo Luciferase Assay Systems kit (Promega) was used for promoter assays according to manufacturer’s protocol. Briefly, ∼2-3 x 10^5^ HEK293 cells were seeded prior to transfection in 12-well plates (Corning Inc.). Cells were transiently transfected with specific pGL3 promoter plasmids (wild type or mutant) in the presence of appropriate vector control or expressing plasmids. 36 h post-transfection cells were harvested, washed with 1 x PBS and suspended with 100 μl of 1 x passive lysis reagent (PLB). 20 μl of the cell extract supernatant was mixed with 100 μl of Luciferase Assay Reagent (LAR) and subjected for luminescence measurement in Synergy H1 microplate reader (BioTek) after a 5 sec delay over a 10 sec window.

### Chromatin immunoprecipitation sequencing (ChIP-Seq)

A total of 2.0 × 10^7^ cells were cross-linked with 1% formaldehyde for 10 min followed by quenching the reaction with 125 mM glycine for 5 min at room temperature. ChIp-Seq was performed using 5 µg E2F1 specific mouse monoclonal antibody (Invitrogen) in two *in vitro* EBV transformed lymphoblastoid cell lines (LCL#1 and LCL#89) using ChIP-IT Express Chromatin Immunoprecipitation Kit (Active Motif, Inc.) according to the manufacturer’s instruction. ChIP libraries were generated using the NEBNext UltraTMII DNA Library preparation kit (New England Biolabs). ChIP Libraries were validated using Qubit 4 fluorometer (Thermo Fisher Scientific) followed by next-generation sequencing analysis on an Illumina HiSeq2500 platform. Reads quality were checked using FastQC followed by adapter trimming with Trimmomatic v0.35. The paired end data aligned to the reference Human (Homo sapiens.GRCh37) using Bowtie2 v 2.5.3 with default parameters. MACS2 call-peak v2.2.9.1 was used for peak calling analysis against the matching input samples. Paired end reads were also aligned with Human gammaherpesvirus 4 genome sequence (RefSeq: NC_007605.1). ChIP-seq signal tracks were visualized using the Integrative Genomics Viewer (https://igv.org/).

### Quantification of EBV genome copy number

Quantification of intracellular and extracellular EBV genome copy number was performed by qRT-PCR analysis. Intracellular viral DNA was extracted from ∼3 × 10^6^ cells by Wizard Genomic DNA Purification Kit (Promega). For extracellular viral DNA isolation, 600 μl cell supernatant was collected and centrifuged at 3000 rcf for 10 min. The supernatant was then treated with 15 μl DNase I (New England Biolabs) at 37^0^C for 30 min followed by heat inactivation at 70^0^C for 10 min. The supernatant was further treated with 15 μl Proteinase K (800 U/ml, New England Biolabs), 100 μl of 10 % (wt/vol) SDS and incubated for 60 min at 65^0^C. DNA was purified using phenol-chloroform method followed by precipitation with sodium acetate and ethanol. Precipitated DNA was dissolved in 50 μl nuclease-free water. The extracted DNA was then diluted to 10 ng/μl and qPCR was performed targeting the EBNA1 viral gene. A standard curve was made by performing qPCR on serial dilutions of Namalwa EBV genome targeting the EBNA1 as previously described (Ryan *et al*, 2004). EBV viral copy number was calculated by putting the sample Cq values into the regression equation provided by the standard curve.

### Lentivirus mediated knockdown (KD) of E2F1 in EBV positive B-cells

Lenti-X 293T cells at ∼70 % confluency in 10-cm cell culture dishes (Corning Inc.) were co-transfected with 10 µg pTRPZ-sh-E2F1 clone along with the two lentivirus packaging plasmids - 4 µg pMDG (Addgene #187440) and 12 µg psPAX2 (Addgene #12260) using CaPO_4_ method as previous described (Malik *et al*, 2024). E2F1 Sh-RNA sequence was adopted from previously published results (Wang *et al*, 2015). 12 h post transfection media was replaced with fresh DMEM with 3 mM NaBu (Merck) to induce lentivirus replication cycle. 48 h post-treatment lentivirus containing media was collected, filtered through 0.45 μM membrane (Corning Inc.) and subjected to spinoculation with ∼5.0 x 10^5^ P3HR1 cells in complete RPMI supplemented with 8 μg/ml polybrene (Merck) at 800 g for 2 h. 48 h post-transduced cells were selected using 1 μg/ml puromycin (Merck) for 7 days. Expression of Sh-RNA in transduced P3HR1 cells was initiated by 1 μg/ml doxycycline (Merck), while transduced cells without doxycycline treatment were served as control. Selected cells with or without doxycycline were harvested and subjected to immunoblot, qRT-PCR, cell proliferation and EBV genome copy number quantification analysis.

### sgRNA CRISPR knockout (KO) analysis

Two individual sets of E2F1 and c-Myc sgRNAs were either designed using the online tool Synthego (https://www.synthego.com/products/bioinformatics/crispr-design-tool) or adopted from previously published manuscripts (Guo *et al*, 2020a; Shats *et al*, 2017). Oligos were obtained from Integrated DNA Technologies, annealed and subsequently cloned into the *Bsm*BI restriction site of the lentiCRISPR v2 vector (Addgene #52961). Lentivirus production followed by spinoculation in P3HR1 cells were performed as described above for stable cell line generation. KO efficiency was validated by immunoblot analysis.

### Cell cycle analysis

Cell cycle analysis was performed as previously described (Malik *et al*, 2024). Briefly, P3HR1 cells were either left untreated or treated with the combination of 20 ng/ml TPA and 3 mM sodium butyrate or specific cell cycle inhibitor 50 μM Mimosine (MedChemExpress) for G0/G1 arrest. 24 h post-treatment cells were washed with 1 x PBS and fixed with ice-cold 70 % ethanol for 30 min at 4^0^C followed by two additional washing steps with 1 x PBS. Fixed cells were treated with staining buffer (5 µg/ml propidium iodide, 40 µg/ml RNase A, 0.1 % Triton X-100 in PBS) for 30 min at room temperature. Each sample was subjected for cell cycle analysis on an S3e Cell Sorter (BIO-RAD). Cell cycle data were analyzed using FCS Express v6.06.0042 (https://denovosoftware.com/).

### Structure prediction and pairwise structure alignment

The predicted structure of E2F1 and E2F2 were acquired using the AlphaFold Protein Structure Database (https://alphafold.ebi.ac.uk/) (Varadi *et al*, 2024). The primary sequence of E2F1 and E2F2 was obtained from UniProt accession IDs - Q01094 and Q14209, respectively. The default parameters of AlphaFold framework were used for the prediction. The confidence of the predicted model was assessed using the per-residue predicted local distance difference test (pLDDT) scores provided by AlphaFold. AlphaFold structure viewer tool was used to perform further structural visualization. Alignment of 3D structures of E2F1 and E2F2 was performed using the RCSB PDB Pairwise Structure Alignment tool.

### Sequence Similarity Consensus analysis

UniProt Align tool (https://www.uniprot.org/align) was used to check the sequence similarity of all E2F transcription factors. Protein sequences of all the E2Fs were extracted from UniProt database (accession IDs: E2F1 - Q01094, E2F2 - Q14209, E2F3 - O00716, E2F4 - Q16254, E2F5 - Q15329, E2F6 - O75461, E2F7 - Q96AV8, E2F8 - A0AVK6) and aligned using default parameters of the UniProt Align tool. Percentage similarity was determined based on the presence of identical residues in the aligned regions. Pairwise alignment and the resulting similarity score was downloaded from the UniProt Align tool for further analysis.

### RNA sequencing and data analysis

P3HR1 cells with or without doxycycline (DOX) containing RPMI for 7 days were either left untreated or treated with EBV lytic cycle inducer - 3 mM NaBU plus 20 ng/ml TPA. 72 h post-induction cells in all four categories were harvested and subjected to RNA isolation. ∼1 µg total RNA was used for library generation using NEBNext Ultra II Directional RNA library Prep Kit (New England Biolabs) followed by RNA sequencing analysis on an Illumina NovaSeq 6000 platform according to the manufacturer’s instructions. For read quality reports FastQC was applied and qualified reads were processed with Trimmomatic v0.35 for trimming the adapter sequences. The sequences were aligned to the Human genome (Homo sapiens.GRCh37) using Bowtie2 v 2.5.3 with default parameters. Gene expression was measured using featureCounts v 2.0.6.

DESeq2 v 2.11.40.8 package from R was utilized to analyse differential expression pattern between experimental groups. Up-regulated and down-regulated genes were selected on the basis of log_2_Fold Change as > = 1.5 and < = −1.5 respectively with p value < = 0.05. Differentially expressed gene sets were further analysed through DAVID v6.8 webserver. Functional analysis was performed by clustering features found across different databases. Gene Ontology (GO) was selected from the hits table for DAVID clustering. The abundance of viral transcripts were quantified utilizing Kallisto quant tool (v. 0.48.0) with Human herpesvirus 4 type 2 reference transcriptome (RefSeq: NC_009334.1) into transcripts per million (TPM), which was further converted to log_2_ (TPM+1) for data representation.

### Statistical analysis

Unless otherwise indicated, all bar and line graphs denote the arithmetic mean of at least three biologically independent experiments (n = 3), with error bars representing standard deviations (SD). Data were analysed using One-Way Anova (Tukey’s multiple comparison test) followed by two tailed student’s t-test or post-Dunnett test to calculate the Statistical significance of differences in the mean values using either GrapPad Prism v8 or Microsoft Excel 2013 software. P-value < 0.05 was considered as significant (*P < 0.05; **P < 0.01; ***P < 0.001; ns, not significant).

## Supplementary Information

**Figure EV1. Differential gene expressions of E2F isoforms during EBV latent infection and lytic cycle reactivation.**

**(A)** Heat map analysis (log2 Fold Change) of all eight E2F isoforms (E2F1-8) of RNA-Seq data (GSE235941) of peripheral blood mononuclear cells (PBMCs) infected with GFP-EBV for 0-4 days post-infection (dpi). **(B)** Heat map representation of reanalysis of microarray data (White *et al*, 2010) of E2F transcripts (E2F1-8) in uninfected and EBV infected BL31 cells. **(C)** Heat map representation of differential gene expression of the E2F isoforms (E2F1-8) of RNA-Seq data (GSE125974) of B-cells infected with EBV for 0-28 dpi. **(D)** qRT-PCR analysis of cDNA generated from PBMCs from two individual donors infected with GFP-EBV for 0-28 DPI. **(E)** Heat map and dot plot analysis of the transcripts profile of the indicated E2F isoforms in whole blood cells (PBLs) and EBV transformed lymphoblastoid cell lines (LCLs) using ‘Genotype-Tissue Expression (GTEx)’ project. **(F)** qRT-PCR analysis of cDNA generated from PBMCs from two individual donors and two LCLs – LCL#1 and LCL#89. **(G)** Heat map representation of differential gene expression of the E2F isoforms (E2F1-8) of RNA-Seq data (GSE237484) of two LCLs (LCL#1 and LCL#89) reactivated to lytic replication by TPA/sodium butyrate (NaBu) for 0-3 days post treatment (dpt). **(H)** Heat map representation of differential gene expression of the E2F isoforms (E2F1-8) of RNA-Seq data (Gain *et al*, 2020) of two LCLs (LCL#1 and LCL#89) either left untreated or treated with 1 µM MG132. qRT-PCR results are presented as the mean ± SD, n = 3 biological replicates. Statistical significance was determined by a two-sided Student’s t-test, *P < 0.05; **P < 0.01; ***P < 0.001; ns, not significant. Source data are available online for this figure.

**Figure EV2. Effect of E2F1 on EBV latent promoters.**

Luciferase reporter activity and the corresponding immunoblot analysis of different EBV latent promoters – Wp, Cp, Qp, LMP1p and LMP2p in the presence of increasing concentrations of E2F1. The results are presented as the mean ± SD, n = 3 biological replicates. Statistical significance was determined by a two-sided Student’s t-test, *P < 0.05; **P < 0.01; ***P < 0.001; ns, not significant. Source data are available online for this figure.

**Figure EV3. Cell cycle arrest does not affect E2F1 expression nor EBV lytic cycle activation.**

**(A-C)** P3HR1 cells either left untreated or treated with Mimosine or TPA/sodium butyrate (NaBu) for 24 h were subjected to either **(A-B)** cell cycle or **(C)** immunoblot analysis. **(D)** Schema showing deletion of pocket protein binding domain (PBD) of E2F1 for cloning in flag-tagged expression vector. **(E)** Immunoblot analysis of HEK293 cells transiently transfected with control vector, flag-tagged wild-type (residues 1-437) E2F1 or pocket protein binding domain deleted E2F1 (residues 1-400) expression plasmids. Cell cycle distribution graphs and blots are representative of n = 3 biological replicates. Source data are available online for this figure.

**Figure EV4. E2F1 knockdown transactivates EBV lytic genes.**

Heatmap analysis of RNA-Seq data of EBV transcripts from P3HR1 cells stably expressing E2F1 sh-RNA in the absence and presence of doxycycline (-/+DOX) and with or without TPA/sodium butyrate (NaBu) treatment for 72 h. Log2 (TPM+1) in EBV mRNA abundance are shown. **(C)** Reanalysis of RNA-Seq data (GSE136597) of all eight E2F isoforms (E2F1-8) in EBV^+^ BL lines Mutu I and Mutu III. **(D)** qRT-PCR analysis of EBV latent and lytic gene mRNAs from P3HR1 cells stably expressing E2F1 sh-RNA in the absence and presence of doxycycline (-/+DOX). qRT-PCR results are presented as the mean ± SD, n = 3 biological replicates. Statistical significance was determined by a two-sided Student’s t-test, *P < 0.05; **P < 0.01; ***P < 0.001; ns, not significant. Source data are available online for this figure.

**Figure EV5. c-Myc does not regulate E2F1 transcription.**

**(A)** Luciferase reporter activity and the corresponding immunoblot analysis of the wild-type c-Myc promoter in the presence of increasing concentrations of E2F1 expression plasmid in transiently transfected HEK293 cells. **(B)** Luciferase reporter activity and the corresponding immunoblot analysis of the c-Myc promoter in the presence of empty vector, wild-type (WT) or transactivation domain deleted (ΔTAD) E2F1 expression plasmids. **(C)** Immunoblot analysis of HEK293T-BAC-GFP-EBV transiently transfected control vector or flag-tagged E2F1 expression plasmid. **(D)** Immunoblot analysis of HEK293T-BAC-GFP-EBV transiently transfected control vector or flag-tagged c-Myc expression plasmid. **(E)** Reanalysis of LCLs ChIP-Seq tracks (GSE30399 and GSE36354) of c-Myc at E2F1 promoter region. **(F)** Luciferase reporter activity and the corresponding immunoblot analysis of the wild-type E2F1 promoter in the presence of increasing concentrations of c-Myc expression plasmid. The results are presented as the mean ± SD, n = 3 biological replicates. Statistical significance was determined by a two-sided Student’s t-test, *P < 0.05; **P < 0.01; ***P < 0.001; ns, not significant. Source data are available online for this figure.

**Figure S1. ChIP-Seq reanalysiqs reveal latent EBV oncoproteins do not occupy E2F1 promoter.**

ChIP-Seq tracks for EBV oncoproteins - EBNA1 (GSE73887), EBNA2 (GSE29498), EBNALP (GSE49338), EBNA3A (GSE88729), EBNA3B (GSE88729), EBNA3C (GSE88729), and NF-κB subunits (GSE55105) - RelA/p65, c-Rel, RelB, p50, p52 at E2F1 promoter region.

**Figure S2. BZLF1 does not affect E2F1 expression at post-translational level.**

**(A)** Immunoblot analysis of HEK293 cells transiently transfected with flag-tagged E2F1 expression plasmid in the presence of either control vector or myc-tagged BZLF1 expression plasmid. **(B)** HEK293 cells transiently transfected myc-tagged BZLF1 expression plasmid with or without flag-tagged E2F1 expression plasmid were subjected to co-immunoprecipitation analysis using anti-flag antibody. Blots are representative of n = 3 biological replicates. Source data are available online for this figure.

**Figure S3. E2F1 and c-Myc transcripts do not positively correlate in EBV negative B-cell lymphomas and solid cancers.**

**(A-B)** Two-sided unpaired Student’s t-test and two-sided Pearson’s correlation were employed to analyse the association between E2F1 and c-Myc transcripts in **(A)** EBV^-^ B-cell, lung cancer, breast cancer, prostate cancer, and melanoma lines from DepMap portal (https://depmap.org/portal/) or **(B)** DLBCL (diffuse large B-cell lymphoma), lung adenocarcinoma (LUAD), breast invasive carcinoma (BRCA), prostate adenocarcinoma (PRAD) and skin cutaneous melanoma (SKCM) patients’ tissue samples from TCGA datasets (https://ualcan.path.uab.edu/). Source data are available online for this figure.

**Figure S4. BZLF1 does not regulate c-Myc transcription.**

**(A)** Heat map analysis (log2 Fold Change) of c-Myc transcript of RNA-Seq data (GSE237484) of LCL#1 and P3HR1 reactivated to lytic replication by TPA/sodium butyrate (NaBu) for 0-3 days post treatment (dpt). **(B)** Reanalysis of Raji ChIP-Seq tracks (E-MTAB-7788) of BZLF1 at c-Myc gene locus. **(C)** Luciferase reporter activity and the corresponding immunoblot analysis of the wild-type c-Myc promoter in the presence of increasing concentrations of BZLF1 expression plasmid in transiently transfected HEK293 cells. The results are presented as the mean ± SD, n = 3 biological replicates. Statistical significance was determined by a two-sided Student’s t-test, *P < 0.05; **P < 0.01; ***P < 0.001; ns, not significant. Source data are available online for this figure.

## Data availability

E2F1 ChIP-Seq datasets in two EBV transformed lymphoblastoid cell lines – LCL#1 and LCL#89, and RNA-Seq datasets for E2F1 knockdown in P3HR1 cells in the absence and presence of EBV lytic cycle inducer TPA/NaBu have been submitted to GEO repository with accession number GSE284473 and GSE285456, respectively. All data generated and analysed in this study are included in the manuscript and supporting files. The raw data for each figure and the numerical data for this study are provided with the manuscript. Correspondence and material requests should be directed to Dr. Abhik Saha. Expanded view data, supplementary information, appendices are available online version for this paper.

## Author contributions

JB designed the study and performed the experiments. SKAA performed part of the experiments. SM involved in data curation and formal analysis. SN involved in data curation. PM revised the manuscript. AS designed the study and wrote the manuscript.

## Acknowledgements

We sincerely thank to Erle S Robertson (Perelman School of Medicine, University of Pennsylavania, USA), Rupak Dutta (Indian Institute of Science Education and Research, Kolkata, India), Debanjan Mukhopadhyay (Presidency University, Kolkata), and National Centre for Cell Science (NCCS), Dept. of Biotechnology (DBT), Govt. of India for providing reagents, plasmids, and cell lines. This work was supported by grants from Department of Biotechnology (DBT), Govt of India (BT/PR40894/MED/29/1532/2020), DBT/Wellcome Trust India Alliance (IA/I/14/2/501537) and Science and Engineering Research Board (SERB) under Department of Science and Technology (DST), Govt. of India (CRG/2023/004657) to AS. The funders had no role in study design, data collection and analysis, decision to publish, or preparation of the manuscript. JB, SM, SN are the recipients of UGC-NET Research Fellowship, India.

## Supplementary Material

**Source Data for Figure 1**

**Source Data for Figure 2**

**Source Data for Figure 3**

**Source Data for Figure 4**

**Source Data for Figure 5**

**Source Data for Figure 6**

**Source Data for Figure 7**

**Source Data for Figure 8**

**Expanded View Figures**

**Source Data for Expanded View Figure 1**

**Source Data for Expanded View Figure 2**

**Source Data for Expanded View Figure 3**

**Source Data for Expanded View Figure 4**

**Source Data for Expanded View Figure 5**

**Table EV1**

**Table EV2**

**Table EV3**

**Dataset EV1**

**Dataset EV2**

## Appendix PDF

**Appendix Figure S1**

**Appendix Figure S2**

**Appendix Figure S3**

**Appendix Figure S4**

